# Deconstructing replicative senescence heterogeneity of human mesenchymal stem cells at single cell resolution reveals therapeutically targetable senescent cell sub-populations

**DOI:** 10.1101/2022.01.24.476823

**Authors:** Atefeh Taherian Fard, Hannah Leeson, Julio Aguado Perez, Giovanni Pietrogrande, Dominique Power, Cecilia Liliana Gomez Inclan, Huiwen Zheng, Christopher Nelson, Farhad Soheilmoghaddam, Nick Glass, Malindrie Dharmaratne, Ebony R. Watson, Jennifer Lu, Sally Martin, Hilda Pickett, Justin Cooper-White, Ernst Wolvetang, Jessica C. Mar

## Abstract

Cellular senescence is characterised by a state of permanent cell cycle arrest. It is accompanied by often variable release of the so-called senescence-associated secretory phenotype (SASP) factors, and occurs in response to a variety of triggers such as persistent DNA damage, telomere dysfunction, or oncogene activation. While cellular senescence is a recognised driver of organismal ageing, the extent of heterogeneity within and between different senescent cell populations remains largely unclear. Elucidating the drivers and extent of variability in cellular senescence states is important for discovering novel targeted seno-therapeutics and for overcoming cell expansion constraints in the cell therapy industry. Here we combine cell biological and single cell RNA-sequencing approaches to investigate heterogeneity of replicative senescence in human ESC-derived mesenchymal stem cells (esMSCs) as MSCs are the cell type of choice for the majority of current stem cell therapies and senescence of MSC is a recognized driver of organismal ageing. Our data identify three senescent subpopulations in the senescing esMSC population that differ in SASP, oncogene expression, and escape from senescence. Uncovering and defining this heterogeneity of senescence states in cultured human esMSCs allowed us to identify potential drug targets that may delay the emergence of senescent MSCs *in vitro* and perhaps *in vivo* in the future.

## Introduction

Cellular senescence is one the main drivers of ageing and age-associated diseases [1]. A variety of cell stressors such as telomere attrition, oncogene activation, and genotoxic or oxidative stress can induce senescence [2]. Depending on the cell type and cellular context, senescent cells will secrete a variety of pro-inflammatory cytokines, chemokines, growth modulators, proteases and other soluble signalling factors, collectively termed senescence-associated secretory-phenotype (SASP) factors [3]. These SASP factors promote inflammation and senescence in neighbouring cells, thus furthering the ageing process itself and accelerating the onset of age-related diseases [3]. Senescence of mesenchymal stem cells (MSCs) is thought to be a particularly important contributor to organismal ageing. This is perhaps most pertinently shown by the devastating accelerated ageing syndromes such as Progeria and Werner syndrome (WS) that exhibit pathologies mainly associated with degeneration of mesenchymal tissues [4]. Encouragingly, transplantation of healthy MSCs into a WS mouse model improved both mean life span and bone density [5].

MSCs are self-renewing multipotent immune-modulatory cells that often home to sites of injury, differentiate into bone, cartilage, fat and smooth muscle cells. MSCs are also an important component of the niches of hematopoietic and other stem cells and play important roles in supporting the vascular system. Hundreds of clinical trials involving autologous and heterologous transplantation of bone marrow, or adipose tissue - or human embryonic stem cells (hESCs) derived MSC are ongoing, particularly for age-related diseases such as osteoporosis and osteoarthritis that are accompanied by an age-dependent loss of both MSC number and potency. To reach clinically relevant cell numbers, MSC populations often need to be expanded for prolonged periods of time through *in vitro* culture expansion. Perhaps not unexpectedly, such *in vitro* cultured MSCs acquire a senescent phenotype over extended passages [6], and this occurs even earlier in primary MSCs derived from donors of advanced age [7]. Previous research has shown that MSC senescence programs not only elicit permanent cell cycle arrest, but also alter the differentiation propensity of MSC from bone or cartilage in the young to increased adipogenesis with advanced age, and results in an erosion of their immuno-modulatory properties, further limiting their clinical efficacy [8]. Previous studies have further shown that senescence of cultured murine and human MSCs is associated with telomere shortening, and is modulated through both p53 and p16/pRb tumor suppressor pathways, as well as GATA4 regulated pathways. Bulk MSC cultures also exhibit notable changes in chromatin organisation and gene expression as these cells undergo replicative senescence [9].

While these studies have certainly started to provide insights into the senescence programs of MSCs, most overlook the fact that MSC cultures represent a highly heterogeneous population of cells composed of stem cell and progenitor compartments that differ in the abundance and distribution of MSC markers [10]. The systematic investigation of MSC senescence is further hindered by the fact that the propensity to enter into senescence is affected by the source of the MSCs [11–21], the stage of the cell differentiation process [18, 19, 22] cultivation times [23–28], medium composition, and donor age. Furthermore, while senescent MSCs are generally identified based on the combined presence of multiple biochemical markers such as expression of p16, p53, p21 and senescence-associated β-galactosidase (SA-β-Gal) staining, previous studies (and our data presented here) clearly show that the levels of these markers are not consistently and homogeneously distributed amongst individual cell sub-populations and senescent cells [29].

To minimize such confounding factors, we have used MSC generated from Schwann cell precursors derived from a single hESC line, to specifically eliminate the impacts of inter-donor variability, source, and donor age (as hESCs are epigenetically reset to a fetal stage). We examined the temporal acquisition, levels and distribution of senescence markers and gene expression in individual cells over the course of senescence acquisition using automated image analysis and single cell RNA sequencing. Our data reveal that subsets of MSCs differentially enter heterogeneous pre-senescent states over time in culture and then progress into two different senescent cell states and a population that has escaped senescence. We identified genes *NUPR1*, *GDF15*, *S100A6*, *PCSK1N*, *MT2A*, *IGFBP5*, *CKS2* and *BIRC5* as important drivers of these different senescent states and predict that interference with the expression of these genes may be able to destabilize these specific MSC senescent sub-states.

## Materials and Methods

### MSC differentiation and culture

hESC line Genea022 [30] was maintained under feeder-free conditions on extracellular matrix (ECM, Sigma) coated plates with mTeSR^™^ Plus (Stemcell Technologies), with passages were performed using EDTA at 70-80% confluence approximately every 5 days. For the Schwann Cell Precursors (SCPs) differentiation, the hESCs were plated as single cells at a density of 90,000 cells/cm^2^. The next day, the culture medium was switched to SCP differentiation medium: DMEM/F12 supplemented with 1X B-27 without Vitamin A, 1X GlutaMax, 1X NEAAs, 1X PenStrep, 100 μM β-mercaptoethanol, 10 μg/mL Holo-transferrin, 10 ng/mL Heregulin B, 3 μM CHIR, 10 μM SB431542, 50 μg/mL ascorbic acid, and 8 ng/mL bFGF. Cells were passaged every 4-5 day with Accutase and maintained in differentiation medium for 21 days. Differentiation to MSCs was initiated by exchanging SCP medium to DMEM low glucose (1 g/L) supplemented with 10% FBS and 1X PenStrep. Date of exchange to MSC medium was designated MSC differentiation day 0. Subsequently, MSCs were passaged using TrypLE at 80% confluence approximately every 5 days and plated in tissue-culture treated flasks. Time between passages increased with increasing senescence levels. For proliferation experiments, MSC differentiation replicates 1, 2 and 3 frozen at the different timepoints T=0, T=1 and T=2 were revived and left to recover overnight. The day after cells were detached, counted, and plated in quintuplicate in a 96 multiwell plate (PerkinElmer) at a confluence of 5000 cells/well. The following day cells were incubated with BrdU at a final concentration of 20 ug/ml for 24 or 6 hours. At the end of the incubation period cells were washed with PBS twice and fixed with PFA 4% for 10 minutes at room temperature.

### Senescence-associated β-galactosidase assay (SA-β-Gal)

Cells were washed in PBS, fixed for 10 minutes in 4% PFA, washed, and incubated at 37°C (in the absence of carbon dioxide) with fresh SA-β-Gal stain solution (pH 6.0): Potassium ferricyanide 5 mM, Potassium ferrocyanide 5 mM, Sodium dihydrogen phosphate 0.4 M, Sodium hydrogen phosphate 92 mM, Sodium chloride 150 mM, Magnesium dichloride 2mM and 1 mg ml^-1^ of 5-bromo-4-chloro-3-indolyl-β-D-galactopyranoside. Staining was evident in 2-4 hours and maximal in 12-16 hours.

### Western Blot

Cells were lysed with RIPA buffer containing protease and phosphatase inhibitors, and samples were prepared at 30 μg of protein with DTT (100 mM) and 1X Laemmli SDS loading dye. Lysates were resolved using denaturing TGS (Tris/glycine/SDS) buffer-based polyacrylamide gel electrophoresis (SDS-PAGE) followed by wet transfer (Tris/glycine/methanol) to nitrocellulose membranes. Primary antibodies Sox10 (rabbit, Cell Signaling Technologies [CST] #89356, 1:1000), p21 (rabbit, CST #2947, 1:1000) and B-actin (mouse, CST # 3700, 1:5000) were incubated at 4°C overnight, and HRP-conjugated secondary antibodies for one hour at room temperature. Cross-reactivity was detected using Clarity ECL (BioRad), and captured images were analyzed using Image Lab 4.1 (Bio-Rad, USA) software.

### Immunochemistry

MSCs were plated in 96 well imaging plates (Costar) coated with ECM and allowed to adhere overnight. Cells were washed once with PBS prior to 10 min fixation with cold 4% paraformaldehyde. Cells were permeabilised with Triton-X100 at 0.1% and blocked with 3% BSA for 1 hour. Primary antibodies (anti-p21 (Cell Signalling, 2946S, 1:400), anti-BrdU (Abcam, AB6326-100UG, 1:400) and anti-p16 (Abcam, AB108349-100UL, 1:400) were incubated overnight at 4°C. Secondary antibodies anti-Mouse IgG (Invitrogen, A11029, 1:400), anti-rat IgG (Abcam, AB150167-500UG, 1:400) and anti-rabbit IgG (Invitrogen, A10042, 1:400) were incubated for 45 minutes at room temperature. Nuclei were counterstained with Hoechst 33342 (Invitrogen, H3570, 2 μg ml^-1^) or 4’,6-diamidino-2-phenylindole (Thermo Scientific, 62248, 1 μg ml^-1^) prior to imaging.

Fluorescence images were acquired using an Operetta CLS High-Content Analysis System with a 10x objective. All the images were analysed using the same pipeline for experimental and biological replicates in the CellProfiler software. The individual cells were identified using the Hoechst counterstaining. Hoescht is a cell permeable dye that intercalates DNA, thus we used the intensity of the signal to estimate the content of genomic which is indicated later as ‘nuclear intensity’. Other nuclear characteristics like size and circularity were also analysed. All the immunofluorescence signals within the nuclei were considered for analysis along with the extra-nuclear signal of the SA-β-Gal activity, which can be detected in the 647 channel.

### Correlation analysis for molecular marker intensities

We used the interquartile range (IQR) method to identify and remove outliers in the intensity data. A data-point was identified as an outlier if it was above the 75^th^ or below the 25^th^ percentile by a factor of 1.5 times the IQR. The data was scaled by dividing each intensity value by the maximum intensity for each marker across time points. Wilcoxon ranked test (p-value < 0.05) was used to measure the difference between marker intensities. Pearson’s correlation was used to compute the correlation between marker intensities. The bimodality index was computed using [31]. An index value of > 1.1 is considered as a “true” bimodal pattern. To identify within cluster correlations, a two-component mixture model [32] was applied on each correlation pair across time points and calculated the correlation within each cluster.

### Telomere Dysfuction Induced Foci (TIF) analysis

MSC cells were grown on Alcian blue coated coverslips for 24 hrs. The following day the coverslips were rinsed in PBS and then fixed for 10 min in freshly prepared 4% paraformaldehyde. Cell permeabilization was performed for 10 min using KCM buffer (120 mM KCl, 20 mM NaCl, 10 mM Tris-HCL pH 7.5, 0.1% Triton X-100). Coverslips were blocked with antibody-dilution buffer (20 mM Tris–HCl pH 7.5, 2% (w/v) BSA, 0.2% (v/v) fish gelatin, 150 mM NaCl, 0.1% (v/v) Triton X-100 and 0.1% (w/v) sodium azide) for 1 hr at room temperature followed by incubation with a ɣH2AX antibody (05-636 Sigma-Aldrich, Anti-phospho-Histone H2A.X (Ser139) Antibody, clone JBW301) overnight at 4°C. The next day, 3×10 min PBS washes were performed, coverslips were incubated with a fluorophore-conjugated secondary antibody for 1hr at room temperature, followed by three more PBS washes. Next, coverslips were fixed again with 4% PFA for 15 min at room temperature. Cells were then dehydrated with an ice-cold ethanol series of 70%, 80%, 90%, dried, and incubated with a TAMRA–OO-(CCCTAA)3 telomeric PNA probe (Panagene) prepared at 0.3 μg/ml in PNA hybridization solution (70% deionized formamide, 0.25% (v/v) NEN blocking reagent (PerkinElmer), 10 mM Tris–HCl, pH 7.5, 4 mM Na2HPO4, 0.5 mM citric acid, and 1.25 mM MgCl2) for 10 min at 80°C. Hybridization was then allowed to occur overnight at room temperature in a humidified chamber. The following day, coverslips were washed for 5 min each in 50% deionized formamide in 2X SSC, 2X SSC, and 2X SSC + 0.1% Tween 20, at 43°C. Finally, cells were counterstained with DAPI and mounted in ProLong^™^ gold antifade reagent. Microscopy images were acquired on a Zeiss Axio Imager microscope with appropriate filter sets. Images were analysed for telomere intensity, ɣH2AX foci, and telomeres colocalising with ɣH2AX using Cellprofiler v2.1.1 [33].

### Library preparation and scRNA-sequencing

Cells were harvested by TrypLE and dead cells were stained with propidium iodide (PI). Live cell FACS was used to collect a healthy population of single cells for single cell RNA sequencing.

Single cell suspensions were sorted by FACS, spun down to concentrate and a cell count was performed to determine post-sort viability and cell concentration (concentration range 7.40E+05 – 2.34E+06, viability 85-94%). Single cell suspension was partitioned and barcoded using the 10X Genomics Chromium Controller (10X Genomics) and the Single Cell 3’ Library and Gel Bead Kit (V2; 10X Genomics; PN-120237). The cells were loaded onto the Chromium Single Cell Chip A (10X Genomics; PN-120236) to target 10,000 cells. GEM generation and barcoding, cDNA amplification, and library construction was performed according to the 10X Genomics Chromium User Guide. Reactions were performed in a C1000 Touch thermal cycler with a Deep Well Reaction Module (Bio-Rad). 11 cDNA amplification cycles were performed, and half of the cDNA was used as input for library construction. 10-13 SI-PCR cycles were used depending on amount of input cDNA. The resulting single cell transcriptome libraries contained unique sample indices for each sample. The libraries were quantified on the Agilent BioAnalyzer 2100 using the High Sensitivity DNA Kit (Agilent, 5067-4626). Libraries were pooled in equimolar ratios, and the pool was quantified by qPCR using the KAPA Library Quantification Kit - illumina/Universal (KAPA Biosystems, KK4824) in combination with the Life Technologies Viia 7 real time PCR instrument. After the initial sequencing run, libraries were re-pooled according to estimated captured cells as determined using the Cell Ranger software (10X Genomics).

Library preparation was performed at the Institute for Molecular Bioscience Sequencing Facility (University of Queensland). Denatured libraries were loaded onto an Illumina NextSeq-500 and sequenced using a 150-cycle High-Output Kit as follows: 26bp (Read1), 8bp (i7 index), 98bp (Read2). Read1 supplies the cell barcode and UMI, i7 the sample index, and Read2 the 3’ sequence of the transcript. In total, 5 sequencing runs were performed.

### scRNA-seq data analysis

#### Pre-processing and quality control

The unfiltered unique molecular identifier (UMI) count matrix from the nine libraries (three replicates for each of the three time-points) was generated using cellranger (v3.0.2). From sequencing, an average of 22,609 high quality reads were obtained per cell. Reads were mapped to human GRCh38 genome, and an average of 92% reads was mapped confidently to the transcriptome. True cells we identified droplets filtered out through DropletUtils R package. Briefly, this method computes the upper quantile of the top expected barcodes, and orders them based on the library size. Any barcode containing more molecules than the 10% of the upper quantile was considered a cell, and retained for further analysis. Moreover, genes with less than 10% expression across all cells were also filtered out. The scRNA-seq dataset consisted of 22,609 mean reads, 2,527 median genes, and 9,831 median UMI counts per cell (Table S1).

#### Normalisation and integration

Seurat’s [34] integration and SCTransform pipeline was used to integrate the samples from the three replicates and normalise the count matrix using the default parameters. Following correction for batch effect with Seurat’s integration method and normalisation via a regularised negative binomial regression model [34] 86,771 good quality cells were retained for the subsequent analysis. After elimination of genes with extremely low expression (as defined as UMI counts in less than 10% of all cells) 6908 genes were retained (out of a total of 32,838 genes). The object was split by replicate and SCT normalised using 3000 highly variable features.

#### Dimensionality reduction and subgroup identification

Principal component analysis (PCA) was performed in Seurat package. The first 20 PCs were kept based on the eigenvalues, and passed into UMAP for two-dimensional visualisation, with the default parameters.

To identity the number of clusters that best describes the heterogeneity in the single cell population, clustering at 1.2, 0.8, 0.6, 0.5, 0.4 and 0.2 resolution was applied. The result was evaluated and visualised using clustree software package (v0.2.0). Clusters resulted from the 0.5 resolution were selected as the identity classes for cluster specific marker identification.

#### Differential expression analysis

To identify the differentially expressed (DE) genes specific to each time-point and cluster, Wilcoxon Rank Sum test was used. Genes with average expression fold-change of x<-0.25 and x>0.25 (Bonferroni adjusted p-value<0.001) were selected for functional annotation analysis and trajectory inference. Pathway over-representation analysis for each gene set was performed using clusterProfiler [35] against KEGG, hallmark and GO biological pathway signatures. Terms with FDR corrected p-value<0.05 were considered significantly enriched. Gene families were identified using MSigDB database [36] and SASP atlas [37]. Senescence associated genes were compared against CellAge database [38] and the cellular senescence signature from [29]. Oncogene and tumor suppressor genes were identified through COSMIC database (Cancer Gene Census) [39] and TSGene (Tumor Suppressor Gene Database) [40], respectively.

#### Potency evaluation and functional enrichment analysis

The R package SCENT(v1.0.2) was used to quantify the proliferation potency of individual cells [41]. The functional interaction networks were constructed by integrating protein-protein interaction (PPI) network and Spearman-correlation of gene-pairs at each time-point. Genes/proteins were extracted from the built-in reference PPI network (that is obtained by integrating various interaction databases in Pathway Commons [42]) in SCENT; and edges were represented by spearman correlation computer from expression of genes at each time time. Edges with high correlation (*i.e*. ρ > |0.5|) were retained for further analysis. Network modules (densely connected regions) were identified using MCODE app [43] in Cytoscape (v3.8.2). The regulatory interactions between gene pairs were obtained from ReactomeFIViz app in Cystoscape which is connected to the Reactome pathway database [44]. Pathway over-representation analysis for each module was performed through MSigDB [36] against GO Biological process, KEGG pathway database and Hallmark gene sets.

#### Pseudotime analysis and trajectory inference

Slingshot R package (v1.6.0) [45] was used to construct the pseudotime trajectory and scShapes R package (v1.0.0) [46] to identify the differentially distributed genes.

## Results

### esMSC display increased replicative senescence over time in culture

It has previously been shown that hESC derived MSCs (esMSCs) undergo replicative senescence over a similar timeframe as primary BM-derived MSC [47]. We generated esMSC from hESC-derived Schwann cell precursors, and assessed whether these esMSC possess the marker and differentiation profiles of primary MSC, and whether they enter replicative senescence following prolonged *in vitro* culture for 23 (T0), 49 (T1) and 86 (T2) days (Fig. 1a). These specific time points were chosen on preliminary experiments aimed to quantify the percentage of β-gal positive cells after long term culture (T0: <10%, T1: <30%, T2: >70%) (Fig. 1d). As shown in Fig. 1b, at early passage (T0) esMSC exhibit the spindle like morphology and surface marker expression profile of primary MSC (CD105^+^, CD73^+^, CD90^+^, CD34^−^, CD14^−^ and CD19^−^) (Fig. S1). The MSC marker profile did not significantly change between T0, T1 or T2, as indicated by the fact that esMSC from each timepoint met standard requirements for MSC characterisation and expressed Rohart test gene-signatures of MSC identity (Fig. S2) [48]. After 49 days in culture (T1, passage 8 for replicates 2 and 3) cell shape became increasingly irregular and larger (Fig. 1b) and at 89 days (T2, passage 12 for replicates 2 and 3) the vast majority of cells had adopted a large and flattened appearance characteristic of senescent MSCs (Fig. 1b). In agreement with these morphological changes, automated image analysis of >15,000 cells from MSC cultures at each timepoint revealed a progressive temporal increase and shifts in population distribution of senescence markers SA-β-Gal, p16, and p21 (Fig. 1d and Fig. S3). This increase in senescence markers was accompanied by a decrease in BrdU incorporation (Fig. 1d), and a decrease in telomere length over time (Fig. S4a). The adoption of other proposed senescence markers such as SenezRed did not show a significant change across timepoints (Fig. S5), which is in agreement with previous reports that found evidence of telomere erosion but no mitochondrial changes in long-term cultured primary human BM-MSC [49]. Western blot analysis of protein lysates from the esMSC cultures revealed a robust increase in p21 protein expression with increased passage number, and confirmed absence of the Schwann cell marker SOX10 (Fig. 1c). Image analysis revealed that senescent MSC cultures displayed an increase in nuclear size and identified the emergence of two distinct populations at T2 that differ in nuclear staining intensity (Fig. 1d and Fig. S3). Quantification of the number of γH2AX stained cells and telomere damage induced foci (TIFs), both hallmarks of senescence [50], indicated that esMSCs accumulate DNA damage in the form of double strand breaks over time in culture, although this did not translate into significant changes in TIF levels (one-way ANOVA, p-value>0.01) (Fig. S4).

**Figure 1.**
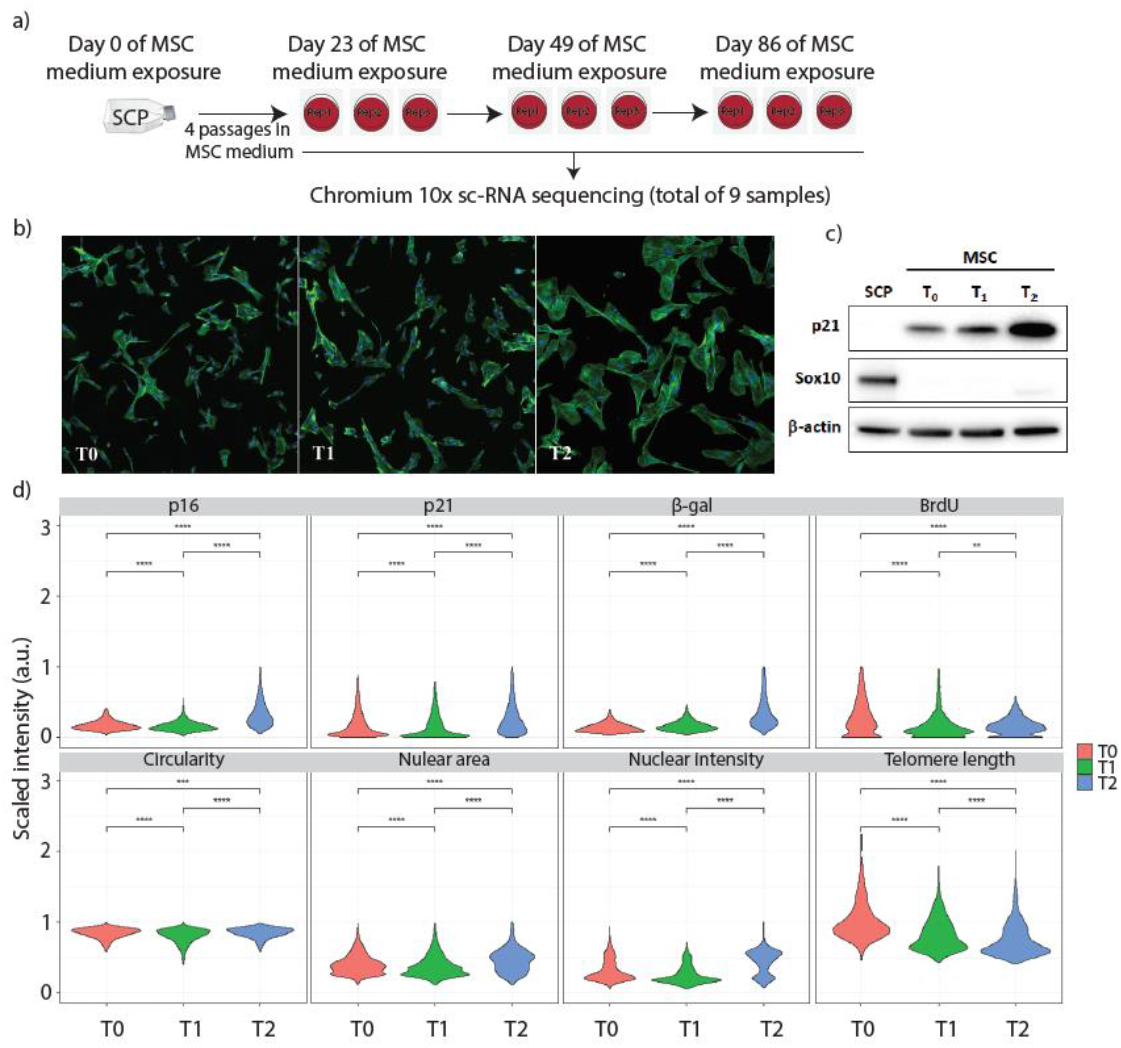
Establishing the human embryonic stem cell (hESC) derive-mesenchymal stem cells (esMSCs) line to study senescence. (a) Schwann Cell Precursors (SCPs) were transdifferentiated into MSCs. Young esMSCs were cultured until population doubling. Day 23, 49 and 86 post MSC medium exposure were selected as T0, T1 and T2 respectively. Cells from each of the three technical replicates at each time-point were harvested for sc-RNA sequencing. (b) Representative images of Young (T0), pre-senescent (T1) and senescent (T2) esMSCs. (c) Western blot analysis to confirm the conversion of SCPs to MSCs by quantifying the abundance of MSC-(p21) and SCP-(SOX10) specific markers. (d) Quantitative estimation of senescence and proliferation specific markers and phenotypic features at T0, T1 and T2 of the experimental set up to confirm the appearance of senescence state. The gradual increase in p16, p21 and β-gal staining are indicative of cell cycle arrest and senescence. Moreover, gradual decrease in Brdu and telomere length also support the presence of the senescence phenotype. There was no substantial difference observed in the circularity feature across different time-points. The bimodal distribution in the nuclear intensity indicates the presence of two cell sub-populations with different DNA contest at T2 (The plots are generated by pooling marker intensity data obtained from replicate 2 and 3; Wilcoxon ranked test, p-value < 0.05).

Given the variability in senescence marker expression distributions across the population over time in culture, we next assessed to what extent the various senescence markers correlated with each other within individual cells. To this end, we computed a Pearson correlation between the intensities obtained from different markers (Fig. 2), and calculated the bimodality index for individual marker intensities (Table S2), revealing that nuclear intensity has the highest bimodality index, specifically at the senescent state, suggesting the presence of two cell subpopulations with differing DNA content. We further noted a strong correlation between nuclear intensity and nuclear area (ρ>0.7) across all three time-points (Fig. S6). By applying clustering on the correlation intensities (Fig. 2) we identified different types of associations between senescence marker intensities in senescent esMSC subpopulations. For example, overall, “nuclear intensity” and “nuclear area” have a positive correlation at T2 (ρ=0.71); but after clustering, the extent of correlation differs in different clusters (cluster 1 ρ = 0.3; cluster 2 ρ = 0.5). Similarly, “nuclear intensity” and “p16” exhibit an overall positive correlation (ρ=0.32), but at the cluster level follows an opposite trend (cluster 1 ρ = −0.35; cluster 2 ρ = 0.3) (Fig. 2). Collectively these data indicated that esMSC undergo replicative senescence, and that there are subsets of cells within the culture that acquire distinct marker profiles suggestive of the presence of multiple senescent states within the esMSC cultures. To further explore this not previously observed heterogeneity in MSC senescence states we next conducted scRNAseq on the same (replicate) cultures that were used for these *in situ* senescence marker analyses.

**Figure 2.**
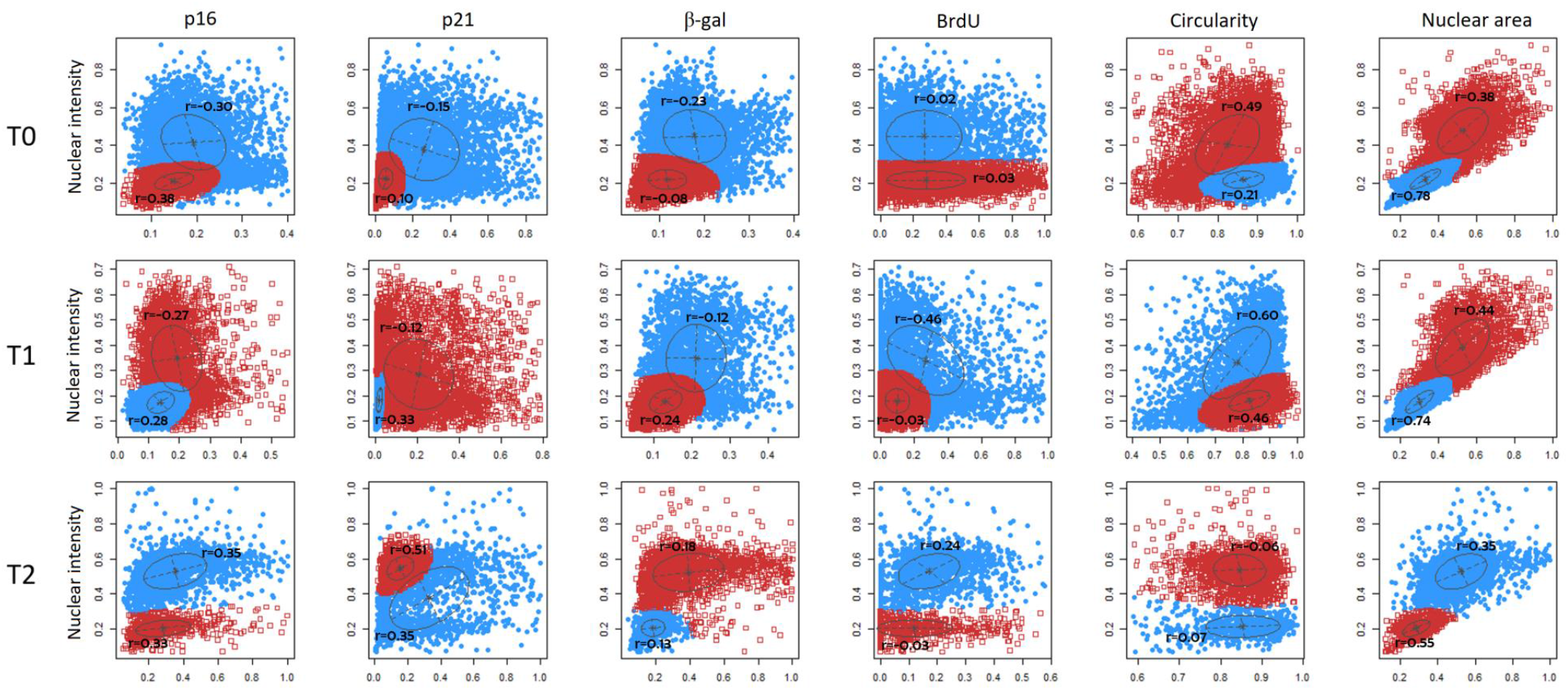
Evaluating the heterogeneity in cell-to-cell associations during senescence. Pearson correlation (represented by r) of nuclear intensity (DNA content) estimates with other molecular markers and phenotypic features at three time points. Clusters (in blue and red) are identified using two-component mixture models. Each cluster represents a subset of cells that show a similar pattern of correlation between each marker pair.

### sc-RNA sequencing of human esMSC undergoing replicative senescence identifies novel senescent states and subpopulations and predicts their hierarchical relationships

Our experimental design consisted of three time points with three biological replicates performed for each time point, resulting in nine separate samples, collectively consisting of 119,454 total sequenced cells with an average of 13,273 cells per sample. Following QC outlined in “Materials and Methods”, we generated a final count matrix comprising of 6908 genes and 86771 high-quality single cells (Table 1). UMAP projection of these data revealed that cells from non-senescent (T0), pre-senescent (T1) and senescent (T2) esMSC clustered separately and were collectively made up of 8 subclusters (Fig. 3a). Indeed, ordering cells along the pseudotime trajectory with Slingshot [45] ordered the cells and cell clusters in a manner consistent to the experimental time course (Fig. 4 and S7). Since the vast majority of cells (>79%) retained expression of MSC cell surface and mRNA markers, as indicated by the fact that 70% of cells passed the Rohart test for MSC identity at each timepoint, these data suggest that the largest contributor to the variation in gene expression between the cell clusters is driven by their step wise progression into pre- and fully senescent cell states over the 86 day time-course. Specifically, Slingshot computed cell trajectories (Fig. 4) indicative of a transition from healthy proliferating esMSCs at T0 (mainly consisting of cluster 2, 6 and a subset of 3), which subsequently transitions into cluster 5, and next cluster 0, the most abundant cell cluster at T1 (52.11%) (Table S3). Cluster 0 cells are next predicted to transition into cluster 4 at T2, which next diverges into two different cell states, cluster 1 and a subset of cluster 3.

**Table 1:** Count matrix dimension before and after pre-processing and filtering.

| Count matrix | Time point (samples) |  |  | Replicate (samples) |  |  |
| --- | --- | --- | --- | --- | --- | --- |
|  | T0 | T1 | T2 | R1 | R2 | R3 |
| Original<br>32838 features x 101945 samples | 31065 | 31065 | 44693 | 43356 | 27200 | 31389 |
| Filtered<br>6908 features x 86771 samples | 27735 | 18389 | 40647 | 33202 | 23975 | 29594 |

**Figure 3.**
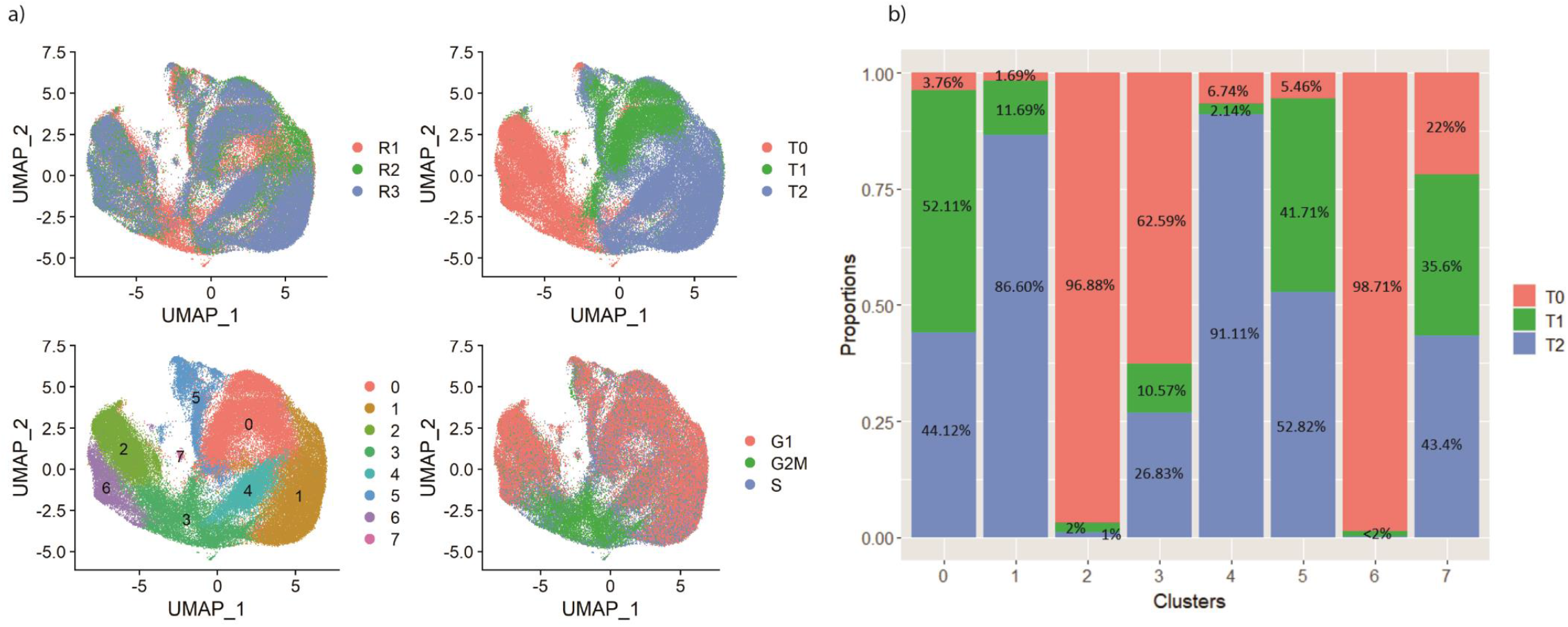
The overall representation of the esMSC population. (a) UMAP of esMSCs color coded by replicate, time-point, cluster and cell cycle phase. (b) Proportion of cells in each cluster with respect to time-points. Cluster 2 and 6 are mainly comprise of T0 cells whereas majority of cells in cluster 1 and 4 are from T2 (senescent) cells. Cluster 0 and 5 comprise of mix of T0, T1 and T2 cells.

**Figure 4.**
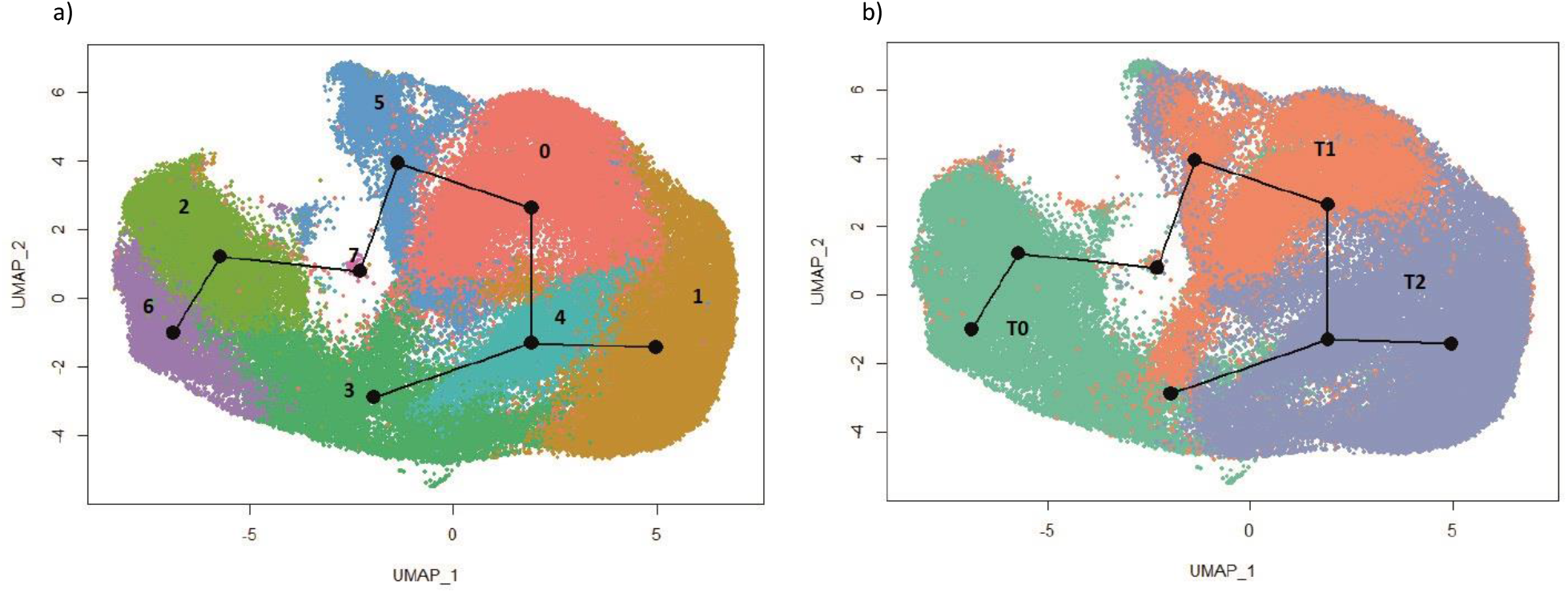
Trajectory inference analysis. The UMAP is color coded based on (a) cluster and (b) time-points. The trajectory starts at cluster 6 and ends at two different senescence states: oncogene related senesce and senescence escapees.

To gain insights into the identity of the computationally identified and temporally ordered esMSC cell clusters that emerge during each of the replicative senescence stages we performed gene enrichment analysis on the top differentially expressed genes of each cluster. This revealed that at T0 cluster 2 displayed significant enrichment for pathways associated with regulation of cell shape, epithelial to mesenchymal transition (EMT) and muscle cell differentiation (Table S4) and displayed high expression of CSRP2, a member of CSRP family encoding a group of short LIM domain proteins that is a critical for cell-cycle progression, development, and smooth muscle differentiation [56] (Tables 2, S4 and S5). In the T0 sub-cluster 3 the top DE genes (Table S6) in esMSC cells were enriched for genes related to regulation of G2/M cell cycle progression. These included UBE2S and PTTG1, genes that are associated with cell-cycle regulation through the anaphase-promoting complex, and CKS1B, which is associated with cell proliferation through regulation of cyclin-dependant protein serine/threonine kinase activity [86]. Cluster 6 of T0 esMSC was enriched for pathways associated with Insulin-like growth factor (IGFs) receptor signalling (Table S4), and exhibited pronounced expression of IGFBP2 and IGFBP4, molecules that are known to play important roles in promoting proliferation, self-renewal, and differentiation of MSCs [57]. Collectively these data indicated that healthy proliferating esMSC consist of at least 3 cell populations that differ in the expression of genes involved in cell cycle progression and differentiation. Cluster 5 contains similar proportions of T1 (41.71%), and T2 (52.82%) cells (Fig. 3b and Table S3). In this cluster, there is enrichment for genes related to metabolic stress regulation of G2/M transition, reactive oxygen species (ROS), oxidative phosphorylation, and Glycolysis (Table S4), pathways previously implicated in metabolic changes that are implicated in promoting senescence [59–61] and ageing [62–65]; higher expression of tissue specific regulators (APOE: Adipogenic, COL1A1: Chondrogenic, THBS1: Osteogenic, ACTA2: Myogenic) and Mitochondrial NADH dehydrogenase subunit genes 2 and 3 (MT-ND2/3) that play an important role in the production of ROS [62]. Indeed, these cells from cluster 5 are computationally projected to transition into cluster 0, which contains roughly equal proportions of T1 cells (52.11%), and T2 cells (44.12%) (Fig. 3b and Table S3). Cluster 0 is enriched for genes related to Apoptosis and p53 pathways (Table S4) and shows strongly increased expression of TRIB3, an inhibitor of cell proliferation [66] that becomes upregulated in response to several forms of cellular stress [67] including oxidative ER stress and hypoxic stress (pathways that are all significantly enriched in cluster 5). We further noted strong expression of *NUPR1* in cluster 0 (Fig. S8 and Table S5), a gene involved in regulating resistance to micro-environmental stress, cell-cycle, apoptosis and DNA repair response [68] and that was previously found to promote K-ras induced senescence in the pancreas of mice [51]. Overall, the results suggest that after undergoing oxidative stress (cluster 5), through stress-induced up-regulation of such genes, MSCs undergo cell cycle arrest, and transition into a pre-senescent state that predominantly consists of cluster 0 cells [69]. T1 cluster 0 cells are transcriptionally close to both cells of cluster 4 and cluster 1. Based on pseudotime trajectory analysis, cluster 0 is the starting point for cells to diverge into different senescent states. Given that the majority of the cells in this cluster are T1, we performed DE analysis of these cells with the other T1 cells in other clusters (pre- and senescent states). These cells show significant high expression of ribosomal protein family, including RPL11 and RPL5 (Fig. S9). In response to oncogenic and replicative stress, the overexpression of these genes mediates p53 activation, which in turn delays proliferation and promotes cellular senescence [52].

**Table 2:** Top 5 time point and cluster-specific markers

|  | Group | No. of cells | No. of markers (up/down) | Top 5 |
| --- | --- | --- | --- | --- |
|  | T0 | 27735 (32%) | 317 (187/130) | KRT18, IGFBP2, SHISA2, MEST, IGFBP4 |
|  | T1 | 18389 (21%) | 126 (77/49) | ACTA2, CRYAB, GDF15, LUM, BGN |
|  | T2 | 40647 (46%) | 240 (83/157) | CCND1, SERPINE2, MMP1, MMP2, NEAT1 |
| Early stage of<br>time course<br>(proliferative) | Cluster2 | 13822 (16%) | 199 (105/94) | KRT18, IGFBP2, FHL1, MAP3K7CL, NREP |
|  | Cluster3 | 11345 (13%) | 360 (267/93) | HIST1H4C, UBE2S, PTTG1, H2AFZ, PCLAF |
|  | Cluster6 | 5229 (6%) | 263 (125/138) | IGFBP2, IGFBP4, IGFBP3, SHISA2, CDH6 |
| Late stage of<br>time course (pre-<br>senescence &<br>senescence) | Cluster0 | 22087 (25%) | 106 (31/75) | NUPR1, BGN, TRIB3, EIF3E, ZFAS1 |
|  | Cluster1 | 19851 (23%) | 200 (60/140) | IGFBP5, MT2A, CCND1, SFRP1, NEAT1 |
|  | Cluster4 | 7595 (8%) | 149 (43/106) | S100A6, G0S2, PCSK1N, CYP1B1, GNG11 |
|  | Cluster5 | 6683 (7%) | 659 (491/168) | POSTN, HSPA5, UBC, HSP90AA1, ACTA2 |
|  | Cluster7 | 159 (<1%) | 940 (137/778) | MALAT1, NEAT1, S100A11, H3F3B, UBA52 |

Cluster 4 is particularly enriched for oncogenes or tumor suppressor genes (23.48%), suggesting these correspond to cells undergoing oncogene associated senescence (Fig. S8, Table S5), including G0-G1 switch gene 2 (*G0S2*), a tumor suppressor gene associated with human dermal fibroblasts senescence, hematopoietic stem cell quiescence, adipocyte differentiation and cell-cycle withdrawal [53]. Another notable gene in cluster 4 is transcription elongation factor A protein-like 7 (*TCEAL7*), a gene that is known to regulate human telomerase reverse transcriptase (*hTERT*) expression and telomerase activity by inhibiting c-Myc pro-oncogene in cells that have activated the alternative lengthening of telomeres (ALT) mechanism. This is significant as more than 70% of mesenchymal tumours use the ALT pathway to maintain the telomere length and bypass replicative senescence [54, 55]. Similar to other clusters, the number of SASP factors are upregulated in cluster 1. These include growth differentiation factor 15 (*GDF15*) [56], *THBS1* [57] and *MMP14* [58] (Fig. S8, Table S5). However, what makes this cluster “SASP-associated” is the exclusive enrichment of TGF-β signalling and inflammatory response pathways, pathways with well-established roles in regulating cellular senescence [58–62]. Notably, there is a strong link between inflammatory response and senescence, as SASP includes inflammatory cytokines and chemokines [58, 63]. The marker genes in cluster 1 that are associated with the inflammatory pathway include *CCL2*, *CD70*, *CDKN1A*, *DCBLD2*, *EREG*, *HIF1A* and *MMP14* (Table S5). Moreover, SASP induces the production and expression of TGF-β, a growth factor known to induce and maintain a senescent phenotype and age-related pathological conditions [59]. We next examined the top significantly up-regulated DE genes in T2 cluster 3 (Fig. 5, Fig. S10 and Table S6). This revealed that these cells show significantly increased expression of SASP factors (*MMP1*, *SERPINE2*, *MMP2*), as compare to the cluster 3 T0 subpopulation (Fig. 5a). They also displayed strong expression of *CCND1*, a well-established regulator of CDK kinases throughout the cell cycle, and a protein that specifically interacts and regulates CDK4/CDK6 that are required for cell cycle G1/S transition [64]. Compared to the rest of T2 cells in the dataset (Fig. 5b and Fig. S10), T2 subcluster 3 displays increased expression of *BIRC5*, an anti-apoptotic gene linked to G2/M cell cycle phase, suggesting that the cells in this cluster have likely escaped cell cycle arrest or never entered cellular senescence in the first place [65]. The fact that this cluster also uniquely over-expresses *TPX2*, a gene that promotes chromosomal instability and escape of cell cycle arrest and senescence [66, 67], further re-inforces the notion that T2 subcluster 3 cells represent putatively oncogenic esMSC that have escaped senescence.

**Figure 5.**
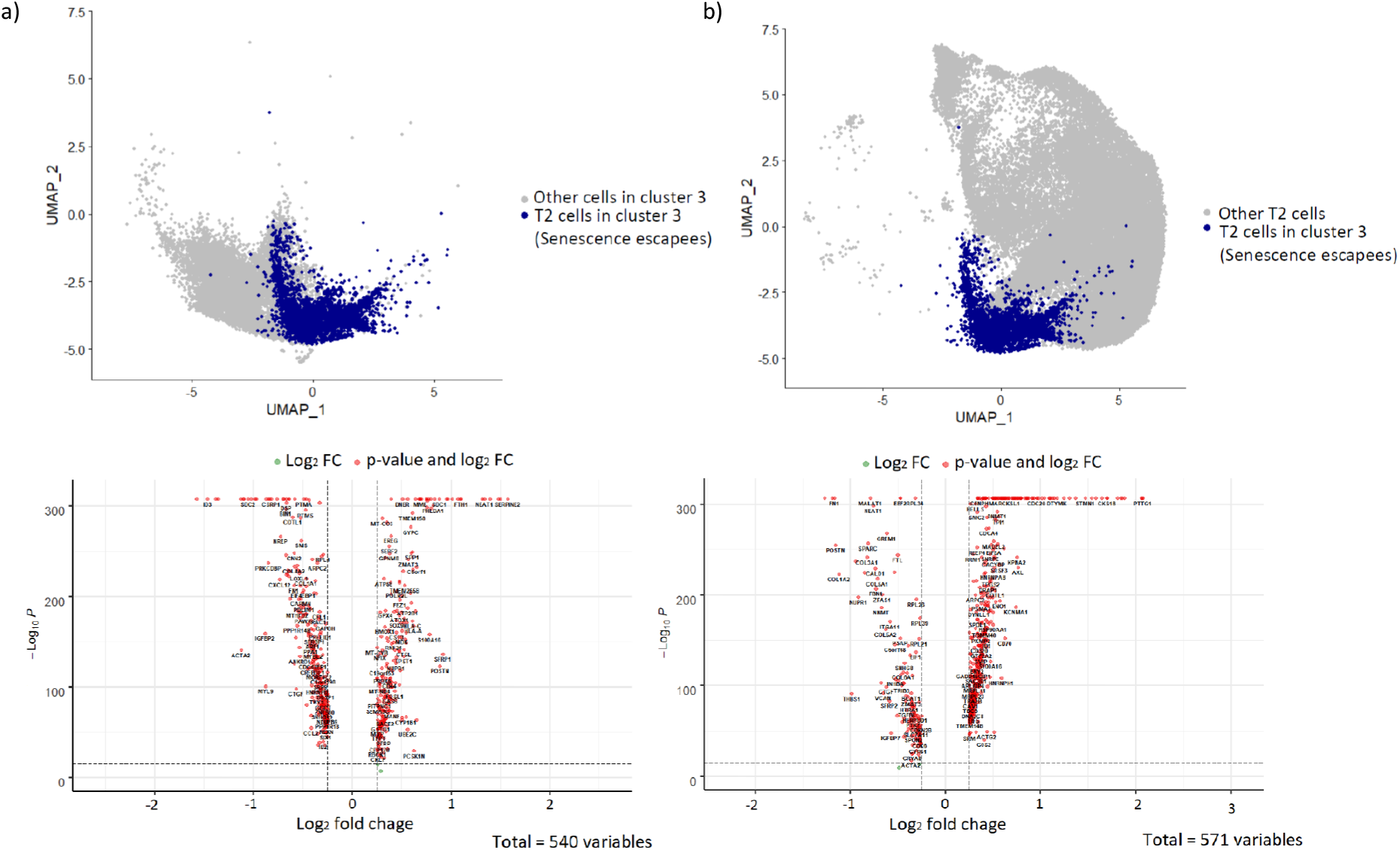
Differential expression analysis of senescence escapees (in blue) with (a) Proliferative esMSCs subtype-3 (other cells in cluster 3), and (b) other senescent cells (other T2 cells). Top panel: UMAP highlight cells in the comparison. Bottom panel: Significant DE genes (LogFC x > |0.25|, Bonferroni adjusted p-value < 0.001)

### Transcription and Gene Regulatory Networks are less tightly regulated in senescent esMSC

Since previous findings have suggested that senescence is associated with an increase in heterogeneity and a loss of robustness in gene expression [68, 69] we next examined transcriptional variability and transcript distribution across the timecourse.

To identify senescence-specific interactions and to study the dynamic behaviour of gene regulatory networks in MSC senescence process, we constructed timepoint-specific gene regulatory networks by integration PPI and gene correlation networks. We extracted the PPI network from the reference database available through SCENT; and computed Spearman correlations of gene expression at each timepoint. Overall, T1 subnetworks displayed the highest number of positively correlated edges, followed by T0 and T2. The genes in subnetworks 4 and 7 are highly positively correlated at T0, but next gradually lose their correlation during senescence. These genes are mainly associated with cell division and regulation of cell cycle transition (*CDC20*, *TPX2*, *CENPF*, *UBE2*, *PPTG1*, *BIRC5* and *CKS1B*). Moreover, in subnetwork 5 only three genes show any sort of correlation in T0 but they develop more positive (and negative) correlation during senescence. The genes in this subnetwork are associated with extracellular matrix organisation (*VCAN*, *SPARC*, *COL1A1* and *COL5A1*) and regulation of tissue development (*COL1A1*, *COL5A1* and *SFRP1*) (Table 3 and S7, Fig. S11). Collectively these data reveal a clear reduction in the number of interactions in both negative and positive correlations as esMSC enter into senescence.

**Table 3:** PPI network comparison during senescence.

| Subnetwork | Nodes | Edges |  |  |  |  |  |
| --- | --- | --- | --- | --- | --- | --- | --- |
|  |  | T0 |  | T1 |  | T2 |  |
|  |  | Positive | Negative | Positive | Negative | Positive | Negative |
| 1 | 31 | 105 | 0 | 450 | 0 | 94 | 0 |
| 2 | 18 | 28 | 15 | 10 | 4 | 5 | 0 |
| 3 | 29 | 29 | 7 | 41 | 0 | 26 | 4 |
| 4 | 8 | 19 | 0 | 0 | 0 | 0 | 0 |
| 5 | 8 | 3 | 0 | 4 | 0 | 9 | 5 |
| 6 | 12 | 6 | 2 | 8 | 2 | 5 | 4 |
| 7 | 5 | 7 | 0 | 1 | 0 | 0 | 0 |
| 8 | 3 | 2 | 0 | 2 | 0 | 1 | 0 |
| Total | 114 | 198 | 24 | 515 | 7 | 140 | 13 |

\*Nodes are extracted from the reference PPI network via SCENT packages and edges represent spearman's $\rho > |0.5|$ . Subnetworks are identified with MCODE app in Cytoscape.

To further examine transcriptional regulation changes across the pseudotime trajectory we next constructed cluster-specific expression networks. The nodes in the networks represent cluster specific marker genes and the edges represent pair-wise Pearson correlation between these genes. Following retention of edges with the highest correlations (*i.e*. ρ > |0.5|) we found that the final networks exhibited a power-law degree distribution, with a few hub genes (Table 4). Highly connected genes in cluster 6 and 2 (T0) are associated with cell proliferation (*e.g*. *IGFBP5*, *TPM1*, *CNN1* and *HBEGF*). Cluster 5 network with the highest number of nodes and edges, parsed into 11 modules, with mitochondrial genes as the top 10 highly connected nodes characterised by genes involved in oxidative phosphorylation and ATP metabolic process (*MT-ATP6*, *MT-CO2*, *MT-CO1*, *MT-ND4*, *MT-ND3*, *MT-CYB*). These observation are consistent with previous data indicating that oxidative stress initiates a DNA damage response that leads to activation of p53 and p16 (CDKN2A) pathways that in turn initiate and sustain cell cycle arrest [70], and supports our inference that cluster 5 (mainly T1 cells) consists of cells undergoing metabolic stress while transitioning from T0 to T2. Moving along the trajectory, cells in cluster 0 (cells transitioning to T2), express genes involved in regulating the recovery from metabolic stress such as TIMP1 and GDF15 that are present in the top 10 most highly connected nodes in cluster 0. TIMP1 is an inhibitor of matrix metalloproteinases (MMPs) and has anti-apoptotic functions [71]. Expression of MMPs is associated with the production of reactive oxygen species (ROS) that drive the emergence of senescence and age-associated disease [72]. GDF15 is involved in the stress response program of cells following cellular injury, and its increased gene expression is associated with age-associated states such as tissue hypoxia, inflammation and oxidative stress [73] and was previously found to contribute to radiation-induced senescence through the ROS-mediated p16 pathway in human endothelial cells [74]. Another gene of note amongst the top 10 highly connected genes in cluster 1 (the senescence state, T2) is *SFRP1*, *a gene* that is commonly over expressed in senescent cells exposed to DNA damage or oxidative stress [75]. Analysis of the co-expression network of cluster 4 of the senescent esMSC indicates that *S100A13* is the gene with the highest number of edges. Previously overexpression of S100A13 was shown to increase NF-κB activity and to induce multiple SASP genes, resulting in the emergence of cellular senescence [76]. Consistent with the notion that T2 cluster 3 is abundant with cells that have escaped senescence we find that the top most highly connected genes in cluster 3 possess both SASP and pro-proliferation properties such as gene involved in G2M checkpoint and cell division.

**Table 4:** Characteristics of cluster-specific co-expression networks.

| Cluster | No. of nodes | No. of edges | No. of modules | Top 10 highly connected genes |
| --- | --- | --- | --- | --- |
| 6 | 115 | 776 | 4 | TKT, UGCG, SLC25A5, AKAP12, EEF1A1, C12orf75, NPM1, HNRNPA1, HBEGF, IGFBP5 |
| 2 | 84 | 546 | 3 | ARPC2, CNN1, PRKCDBP, TPM1, EEF1A1, HMGCS1, IGFBP2, MAP2K2, SELENOW, COL3A1 |
| 5 | 396 | 11932 | 11 | MT-ATP6, VCAN, MT-CO2, MT-CO1, GSTP1, MT-ND4, MT-ND3, FBN1, COL1A1, MT-CYB |
| 0 | 22 | 48 | 1 | TIMP1, COL1A1, RPL21, C6orf48, NPC2, VCAN, TKT, RPS3A, ZFAS1, GDF15 |
| 1 | 42 | 178 | 4 | DCBLD2, S100A16, FTH1, SFRP1, HMGA1, THBS1, CCND1, IGFBP5, MT2A, HMGA2 |
| 4 | 25 | 92 | 2 | S100A13, HLA-A, KCNMA1, CD63, PCSK1N, RPS27, B2M, HLA-B, HMGA1, FTL |
| 3 | 260 | 3552 | 6 | KNSTRN, CENPF, CCNB1, CKS2, ARL6IP1, TPX2, KPNA2, CKS1B, CDC20, BIRC5 |

### Investigating the temporal gene-expression heterogeneity of MSCs during senescence

To identify timepoint specific markers we next assessed the temporal changes in the gene expression levels of esMSCs from T0 to T2 (Fig. 6a). Markers were identified by performing Wilcoxon ranked sum test for cells at each time-point against the rest of the cells. Overall 317, 126 and 240 genes were identified as markers for cells at T0, T1 and T2 respectively (p-value < 0.001; log fold change of > 0.25 up/down-regulated) (Table 2 and Fig. 6b).

**Figure 6.**
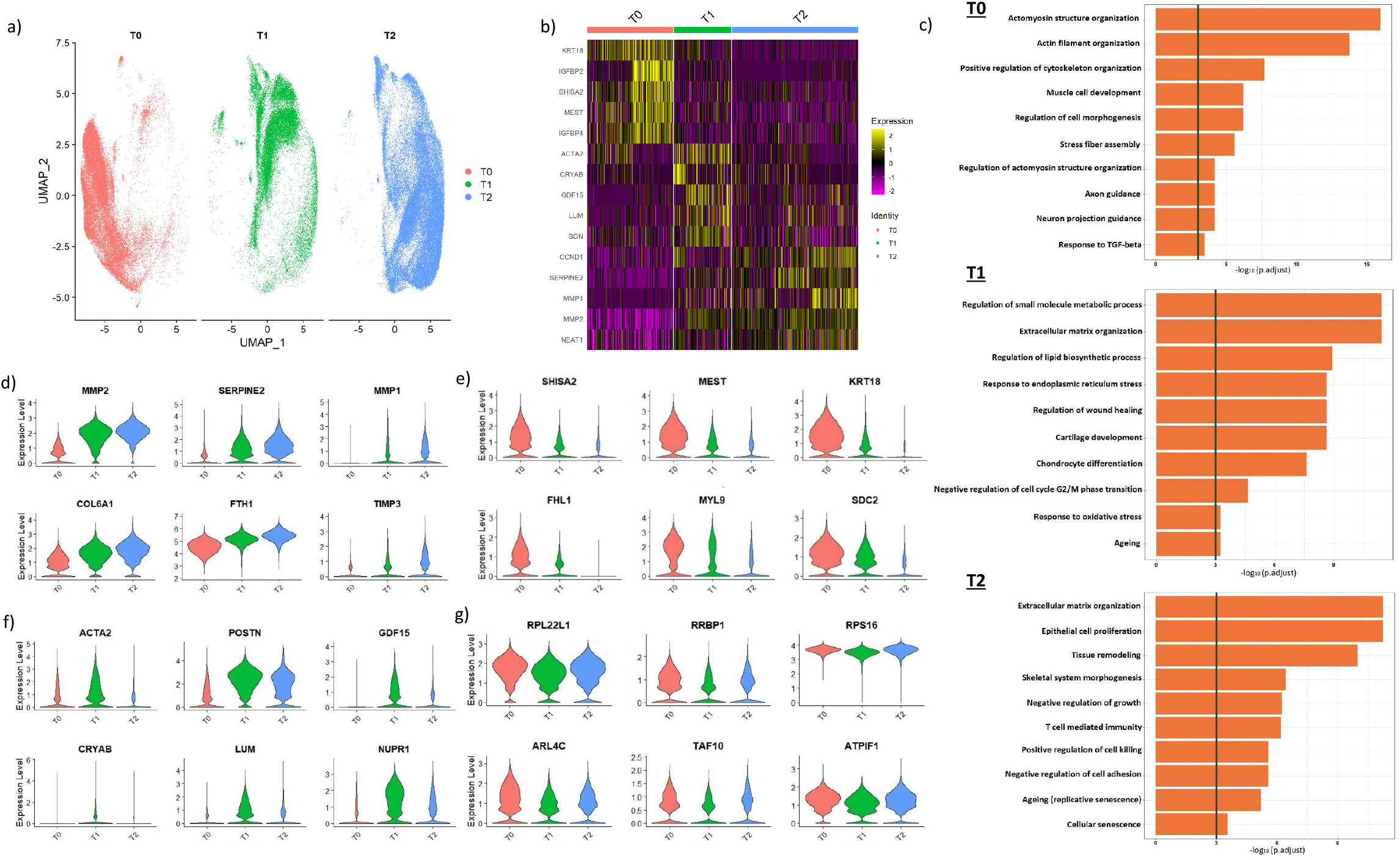
(a) UMAP representation of esMSCs, split by time-point. (b) Normalised gene expression of top five marker genes for each time-point. (c) Pathway over-representation analysis of upregulated genes (logFC > 0.25, p-value < 0.001) in each time-point, against GO database, biological pathways. Pathways with q-value < 0.05 are considered significant. (e-g) Gene clusters with a similar pattern of gene expression during senescence. Differential expression analysis was performed for T0 vs T1 and T1 vs T2. A total of 178 genes had a significant expression change at every time-point. These genes were divided into four groups according to their pattern of gene expression (logFC x> |0.25| Adjusted p.value < 0.001). Genes that consistently (d) increase expression (e) decrease expression (f) transiently increase expression at T1 and (g) transiently decrease expression at T1 during senescence.

At T0, significantly enriched pathways were mainly associated with regulation of cell shape, cellular assembly and organ development (Fig 6c). Response to oxidative stress, angiogenesis and aging pathways were enriched in T1 cells, indicating the tendency of these cells to transition to a stable phenotypic state (whether a specific cell type or a senescent state). Pathways such as ageing, cellular senescence and PI3k-signalling were significantly enriched in the fully senescent population (T2 cells) (Fig. 6c and Table S5).

The senescence-related genes that are the strongest contributors to these temporal changes were identified by assessing differential expression between consecutive time points, *i.e*. T0 *vs*. T1 and T1 *vs*. T2 and identifying clusters of genes within these cohorts that follow the same temporal expression pattern throughout the trajectory. This revealed clusters that either consistently increase or decrease expression over time (Fig. 6d-g and Table S6) or exhibit transient changes (Fig. 6g-h and Table S6). Tissue-specific proliferation genes such as *MEST* (adipocyte differentiation and proliferation [77]) and KRT18 (a member of intermediate filament protein family that provides tissue integrity and structural support in the cytoplasm and nucleus [78]) were among the genes that showed a gradual decrease in expression over time, whereas, SASP factors (e.g. *MMP1* and *SERPINE2*) showed an opposite trend.

We also investigated the expression of the 10 protein-coding mitochondrial genes at the 3 different timepoints (T0, T1, T2) to check whether there was a significant change in their expression during senescence. Except MT-CO2 which displayed a consistently decrease expression, all other genes exhibited a significant increase in expression either at the pre-senescent (T1) or the senescent state (T2) (Fig. S12, Table S8). This observation is consistent with top 10 highly connected genes in cluster-specific co-expression networks (specifically cluster 5, T1 cells), and supports past studies suggesting an increase in production of ROS by dysfunctional mitochondria [79].

### Assessing differential distributions of gene expression reveals genes associated with long non-coding RNA, DNA damage response and apoptosis during replicative senescence

Single cell RNA sequencing data measure cell-specific gene expression, and as a result, provide the opportunity to study the extent of gene expression heterogeneity within a biological condition. Single cell RNA-seq data is complex with high sparsity (increased zeros) and sometimes multimodal gene expression distribution. Therefore, if the data are not modelled reliably, important signals might go unobserved. To capture differences in gene expression heterogeneity beyond simple mean shift, we next investigated gene expression distributions across different timepoints. To this end, gene expression was fitted using generalised linear models with different distributions, including Poison (P), zero inflated Poison (ZIP), negative binomial (NB) and zero inflated negative binomial (ZINB) distributions adjusted for replicate ID. Out of the 3563 that pass the Kolmogorov–Smirnov (KS) goodness of fit test, 585 genes (16.4%) changed distribution in at least two time-points (Tables 5, 6 and S9). Among these, 117 genes (20%) switched distributions at all three time-points that is, from T0 to T1, and from T1 to T2. Pathway over-representation analysis of these genes showed a significant enrichment in genes involved in the response to DNA damage stimuli and non-coding RNA processing pathways (Table S4). These data are in accordance with previous studies that showed lncRNAs that target p21/p53 and pRB/p16 pathways [80] are involved in telomere length attrition [81], consistent with our data showing shortening of telomere length in senescent esMSCs (Fig. S4). Out of the genes that switched distribution at every time-point, genes associated with regulation of cell death and apoptotic process (*RARA*, *KAKRN*, *DLG5*, *FAIM*, *NFATC4* and *APBB2*) changed distribution from unimodal to zero inflated at T1, and back to unimodal distribution at T2. Genes including *CDK3*, *SRGN*, *UHRF1* and *REV3L* changed distribution from zero inflated to unimodal to zero inflated at T0, T1 and T2, respectively (Table 5, 6 & S9). *CDK3* is a cyclin kinase that is an important regulator of cell cycle by promoting G0-G1 and G1-S cell cycle transitions. Its increased expression has been associated with enhanced cell proliferation whereas its knockdown suppresses proliferation in cancer [82]. Similarly, *UHRF1* is essential for maintaining DNA methylation function in a p53-dependent damage checkpoint [83, 84], and its depletion has been associated with G2/M cell cycle arrest, activation of DNA damage response and apoptosis [85].

**Table 5:** Set of genes belonging to the same family of distributions. Total of 3563 genes passed the Kolmogrov-Smirnov test

|  | T0 | T1 | T2 |
| --- | --- | --- | --- |
| Poisson | 1568 (44%) | 1742 (48.89%) | 1556 (43.67%) |
| NB | 1833 (51.44%) | 1647 (46.22%) | 1881 (52.79%) |
| ZIP | 135 (3.78%) | 150 (4.2%) | 106 (2.97%) |
| ZINB | 26 (0.72%) | 23 (0.64%) | 19 (0.53) |

**Table 6:** Differentially distributed genes across time-points.

|  | Count | T0 | T1 | T2 |
| --- | --- | --- | --- | --- |
| Changing in all three time points<br>Total = 117 | 14 | Poisson | NB | ZIP |
|  | 22 | Poisson | ZIP | NB |
|  | 1 | Poisson | ZINB | NB |
|  | 22 | NB | Poisson | ZIP |
|  | 14 | NB | ZIP | Poisson |
|  | 2 | NB | ZINB | ZIP |
|  | 27 | ZIP | Poisson | NB |
|  | 15 | ZIP | NB | Poisson |
| Changing in two time points<br>Total: 468 | 138 | Poisson | NB | Poisson |
|  | 30 | Poisson | ZIP | Poisson |
|  | 212 | NB | Poisson | NB |
|  | 69 | NB | ZIP | NB |
|  | 15 | NB | ZINB | NB |
|  | 1 | ZIP | Poisson | ZIP |
|  | 3 | ZINB | NB | ZINB |
|  | Total 585 |  |  |  |

### A subset of MSCs retain their proliferation potential at the senescent state

To quantify the proliferation potential of MSCs during senescence in an unbiased fashion from transcriptome data we applied signalling entropy rate (SR) analysis, an analysis method that identifies the degree of correlation between transcriptome and connectome (*i.e*. predicted interaction of individual genes with the rest of the PPI network) [86]. SR score range from 1-4, indicating the highest and lowest proliferation potency, respectively (Fig. 7a). Our SR analysis showed that even though esMSC are a heterogeneous cell population with different potency scores at each timepoint, the majority of esMSC lose differentiation potency over time (Fig. 7b) with 43% of cells T2 obtaining the lowest potency score (potency state 4), whereas more than 50% of cells at T0 and T1 had the highest potency score (potency states 1 and 2) (Fig. 7c). As far as the high potency T2 cells are concerned, this observation is in line with the BrdU staining result. From the total number of cells that stained positive for BrdU at T0, 28% remained positive at the T2 stage of experimental design, indicating that a proportion of senescent MSCs have escaped cell cycle arrest associated with senescence, or they have not reached the senescent state yet.

**Figure 7.**
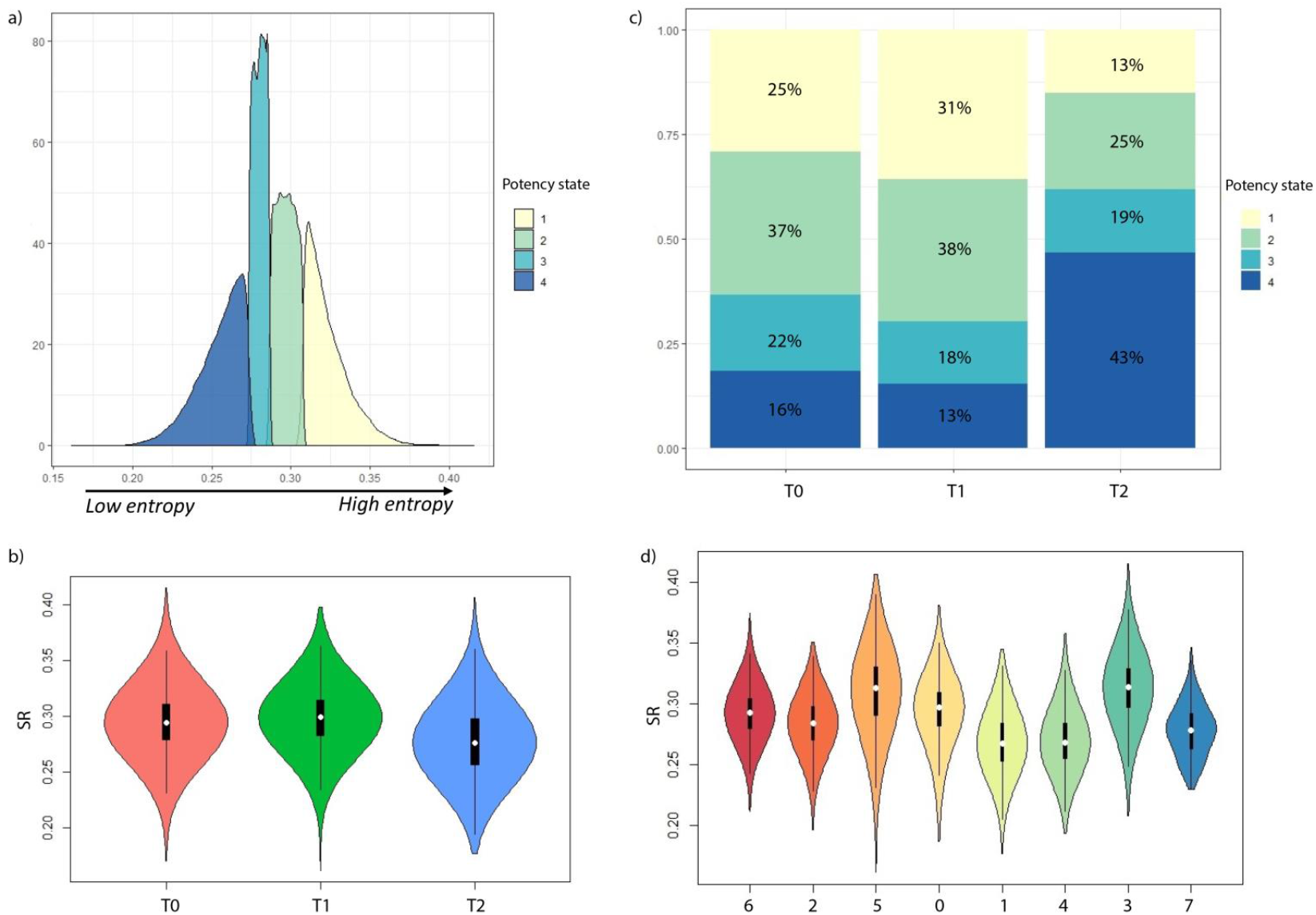
Evaluating the proliferation potential (as a measure of entropy) of esMSCs undergoing replicative senescence. The higher the Shannon entropy rate (SR) the more proliferative the cells are. (a) Based on the SR scores, four potency states were identified. Potency state 1 indicates the highest proliferation potential whereas potency state 4 indicates the least proliferation potential. (b) SR score of cells based on time-point. (c) Proportion of cells with different potency states at each time-point (χ^2^ test, p-value < 0.05). More than 50% of the cells at T0 and T1 are in potency states 1 and 2, whereas majority of cells at T2 acquire a lower potency state. (d) SR score of cells based on cluster.

The SR analysis at the cell subpopulation level showed that cells acquire a higher proliferation potential at T1 (cluster 5), and then gradually lose their proliferation power at the two senescence states (cluster 1 and 4). As expected, Cluster 3 shows a higher variability in SR score since this cluster consist of cells derived from both T0 and T2 (Fig. 7d).

## Discussion

Primary senescence is a cell-intrinsically activated tumour suppressor mechanism brought about by cellular stress. Senescence is considered to be a stochastic process that is driven by both telomere shortening and ensuing telomere damage signalling [87], progressive mitochondrial dysfunction in individual cells, resulting in a heterogeneous occurrence of senescence in progressively increasing numbers of subsets of cells over time in culture as well as *in vivo* [88]. Since primary senescent cells simultaneously induce secondary senescence in neighbouring cells via paracrine and juxtacrine signalling processes that are commonly referred to as SASP, deconstructing the molecular mechanisms of replicative senescence has been challenging and has hampered the design of targeted seno-therapeutics.

To start to address these challenges, here we investigate replicative senescence at a single cell level in hESC-derived MSC, cultured for prolonged periods of time *in vitro*. We first establish that at a population level esMSCs exhibit the marker profile and differentiation properties of primary MSC. We next verify that esMSC display a progressive reduction in BrdU incorporation, an increase in p21 and p16 expression, a reduction in telomere length, an increase in nuclear shape abnormalities and size, an increase in SA-β-Gal staining, elevated SASP factor expression, and characteristic changes in cell shape, as expected of cells undergoing primary and secondary senescence. By measuring these hallmarks of senescence at a single cell level with unbiased high-throughput imaging approaches we identified subsets of cells that acquire distinct marker profiles, suggesting the emergence of multiple senescent states within the esMSC cultures over time in culture, as well as a population of cells that are possibly polyploid and with the potential to escape senescence. Comprehensive single cell transcriptome analysis of the esMSC cultures undergoing progressive replicative and secondary senescence revealed that healthy proliferating, pre-senescent and fully senescent esMSC populations consisted of cell sub-populations bearing distinct transcriptional signatures. Pseudo-temporal analysis of these data sets next showed that healthy “young” passage 1 (day 23) esMSC (clusters 2, 3, and 6) transition at passage 8 into two pre-senescent states that show transcriptional signatures enriched for metabolic stress regulation of G2/M transition, reactive oxygen species (ROS), oxidative phosphorylation, and Glycolysis (cluster 5) and p53 and apoptosis (cluster 0). At passage 12 (day 89) cluster 0 cells transition to cluster 4 that is enriched for oncogene-induced senescence pathways. This population next divides into either cluster 1 cells that show a particularly strong signature of SASP and a population of cells (T2 subcluster 3) that appears to have escaped senescence as indicated by the re-expression of proliferation markers. We deem it highly likely that this population of senescence escapees represents the population of BrdU-labelled polyploid cells observed in our imaging analysis. Overall our data are consistent with the notion that telomere erosion induces a DNA damage response, ROS and mitochondrial dysfunction pathways that activate p53, the retinoblastoma-associated protein (pRb) and the cell cycle-dependent kinase (CDK) inhibitors p16INK4A and p21, that mediate proliferation arrest.

Having established this order of senescent state transitions we next show that the pattern of co-expression is highly heterogeneous at each timepoint, and network modules are specific to each cell subpopulation. We identified a module of highly connected mitochondrial genes in cluster 5 (T1 cells), providing the evidence of oxidative stress response prior to senescence. Our result supports the previous data suggesting mitochondrial genes as potential targets to reduce the effect of senescence [89, 90]. Transcription of a gene is a stochastic process, occurring in irregular bursts or pulses of activity, interspersed by irregular intervals of inactivity [91]. The irregularity in transcriptional bursting is known to be the main driver of diversity of cell behaviours in differentiation and disease, reflecting the underlying mechanisms of transcriptional regulation [92]. We used zero inflation in gene expression distribution as a function of reduced transactional bursting. We found that genes exhibiting zero-inflation in young (T0) and senescent MSCs (T2) were pro-proliferative (*e.g. CDK3*) and pro-apoptotic (*e.g. UHFR1*), respectively. We computed a potency score to assess the proliferation potential of cells during senescence. While there is a mixed population of cells with different levels of proliferation potential in all timepoints, overall, the proliferation potential of cells decreases during senescence.

Collectively our study reveals that replicative senescence of *in vitro* cultured human MSCs results in a temporally ordered sequence of cell state transitions. Our data will enable the design of better culture conditions and intervention strategies that may enable cells to overcome these bottle necks that currently constrain MSC expansion for therapeutic purposes. Our data further permit the identification of senotherapeutics that target the different senescent states that MSC adopt following prolonged culture expansion, and such molecules may also be able to improve MSC ageing *in vivo* in the future.

## Author Contributions

ATF performed computational analysis with input from MD, HZ, ERW, JL. HL, DP, CGI, FS, NG, SM, GP, JAP and CN performed experimental components with assistance from HP, EW, JCW and JCM. Methods sections were contributed by HL, JAP, GP, CGI, CN, HZ and JL. ATF, EW, and JCM reviewed and edited the manuscript. JCM and EW designed the study. All authors read and approved the final manuscript.

## Conflicts of Interest

The authors declare no conflict of interest.

## Notes

### Competing Interest Statement

The authors have declared no competing interest.

## References

1. McHugh, D. and J. Gil, Senescence and aging: Causes, consequences, and therapeutic avenues. The Journal of cell biology, 2018. 217(1): p. 65–77.

2. Childs, B.G., et al., Cellular senescence in aging and age-related disease: from mechanisms to therapy. Nat Med, 2015. 21(12): p. 1424–35.

3. Watanabe, S., et al., Impact of senescence-associated secretory phenotype and its potential as a therapeutic target for senescence-associated diseases. Cancer Sci, 2017. 108(4): p. 563–569.

4. Dreesen, O. and C.L. Stewart, Accelerated aging syndromes, are they relevant to normal human aging? Aging, 2011. 3(9): p. 889–895.

5. Brunauer, R. and B.K. Kennedy, Progeria accelerates adult stem cell aging. Science, 2015. 348(6239): p. 1093–1094.

6. Liu, J., et al., Senescence in Mesenchymal Stem Cells: Functional Alterations, Molecular Mechanisms, and Rejuvenation Strategies. Frontiers in Cell and Developmental Biology, 2020. 8(258).

7. Bruna, F., et al., Regenerative Potential of Mesenchymal Stromal Cells: Age-Related Changes. Stem cells international, 2016. 2016: p. 1461648–1461648.

8. Marędziak, M., et al., The Influence of Aging on the Regenerative Potential of Human Adipose Derived Mesenchymal Stem Cells. Stem cells international, 2016. 2016: p. 2152435–2152435.

9. Zhou, X., et al., Mesenchymal Stem Cell Senescence and Rejuvenation: Current Status and Challenges. Frontiers in Cell and Developmental Biology, 2020. 8(364).

10. Ho, A.D., W. Wagner, and W. Franke, Heterogeneity of mesenchymal stromal cell preparations. Cytotherapy, 2008. 10(4): p. 320–30.

11. Panepucci, R.A., et al., Comparison of Gene Expression of Umbilical Cord Vein and Bone Marrow–Derived Mesenchymal Stem Cells. STEM CELLS, 2004. 22(7): p. 1263–1278.

12. Tsai, M.-S., et al., Functional Network Analysis of the Transcriptomes of Mesenchymal Stem Cells Derived from Amniotic Fluid, Amniotic Membrane, Cord Blood, and Bone Marrow. STEM CELLS, 2007. 25(10): p. 2511–2523.

13. Secco, M., et al., Gene expression profile of mesenchymal stem cells from paired umbilical cord units: cord is different from blood. Stem cell reviews and reports, 2009. 5(4): p. 387–401.

14. Jansen, B.J.H., et al., Functional Differences Between Mesenchymal Stem Cell Populations Are Reflected by Their Transcriptome. Stem Cells and Development, 2009. 19(4): p. 481–490.

15. Wang, T.-H., Y.-S. Lee, and S.-M. Hwang, Transcriptome Analysis of Common Gene Expression in Human Mesenchymal Stem Cells Derived from Four Different Origins, in Mesenchymal Stem Cell Assays and Applications, M. Vemuri, L.G. Chase, and M.S. Rao, Editors. 2011, Humana Press: Totowa, NJ. p. 405–417.

16. Kim, S.-H., et al., Gene expression profile in mesenchymal stem cells derived from dental tissues and bone marrow. Journal of periodontal & implant science, 2011. 41(4): p. 192–200.

17. Miranda, H.C., et al., A quantitative proteomic and transcriptomic comparison of human mesenchymal stem cells from bone marrow and umbilical cord vein. PROTEOMICS, 2012. 12(17): p. 2607–2617.

18. Roobrouck, V.D., et al., Differentiation Potential of Human Postnatal Mesenchymal Stem Cells, Mesoangioblasts, and Multipotent Adult Progenitor Cells Reflected in Their Transcriptome and Partially Influenced by the Culture Conditions. STEM CELLS, 2011. 29(5): p. 871–882.

19. Yoo, H.J., et al., Gene expression profile during chondrogenesis in human bone marrow derived mesenchymal stem cells using a cDNA microarray. Journal of Korean medical science, 2011. 26(7): p. 851–858.

20. Medeiros Tavares Marques, J.C., et al., Identification of new genes associated to senescent and tumorigenic phenotypes in mesenchymal stem cells. Scientific Reports, 2017. 7(1): p. 17837.

21. Kornicka, K., J. Houston, and K. Marycz, Dysfunction of Mesenchymal Stem Cells Isolated from Metabolic Syndrome and Type 2 Diabetic Patients as Result of Oxidative Stress and Autophagy may Limit Their Potential Therapeutic Use. Stem cell reviews and reports, 2018. 14(3): p. 337–345.

22. Menssen, A., et al., Differential gene expression profiling of human bone marrow-derived mesenchymal stem cells during adipogenic development. BMC Genomics, 2011. 12(1): p. 461.

23. Izadpanah, R., et al., Long-term <em>In vitro</em> Expansion Alters the Biology of Adult Mesenchymal Stem Cells. Cancer Research, 2008. 68(11): p. 4229–4238.

24. Ryu, E., et al., Identification of senescence-associated genes in human bone marrow mesenchymal stem cells. Biochemical and Biophysical Research Communications, 2008. 371(3): p. 431–436.

25. Ning, X., et al., Changes of biological characteristics and gene expression profile of umbilical cord mesenchymal stem cells during senescence in culture. Zhongguo shi yan xue ye xue za zhi, 2012. 20(2): p. 458–465.

26. Yoo, J.K., S.-j. Choi, and J.K. Kim, Expression profiles of subtracted mRNAs during cellular senescence in human mesenchymal stem cells derived from bone marrow. Experimental Gerontology, 2013. 48(5): p. 464–471.

27. Ren, J., et al., Intra-subject variability in human bone marrow stromal cell (BMSC) replicative senescence: Molecular changes associated with BMSC senescence. Stem Cell Research, 2013. 11(3): p. 1060–1073.

28. Stolzing, A. and A. Scutt, Age-related impairment of mesenchymal progenitor cell function. Aging Cell, 2006. 5(3): p. 213–224.

29. Casella, G., et al., Transcriptome signature of cellular senescence. Nucleic Acids Research, 2019. 47(14): p. 7294–7305.

30. Dumevska, B., et al., Derivation of human embryonic stem cell line Genea022. Stem Cell Research, 2016. 16(2): p. 472–475.

31. Wang, J., et al., The bimodality index: a criterion for discovering and ranking bimodal signatures from cancer gene expression profiling data. Cancer informatics, 2009. 7: p. 199–216.

32. Scrucca, L., et al., mclust 5: Clustering, Classification and Density Estimation Using Gaussian Finite Mixture Models. The R journal, 2016. 8(1): p. 289–317.

33. Carpenter, A.E., et al., CellProfiler: image analysis software for identifying and quantifying cell phenotypes. Genome Biology, 2006. 7(10): p. R100.

34. Hafemeister, C. and R. Satija, Normalization and variance stabilization of single-cell RNA-seq data using regularized negative binomial regression. Genome Biology, 2019. 20(1): p. 296.

35. Yu, G., et al., clusterProfiler: an R Package for Comparing Biological Themes Among Gene Clusters. OMICS: A Journal of Integrative Biology, 2012. 16(5): p. 284–287.

36. Liberzon, A., et al., Molecular signatures database (MSigDB) 3.0. Bioinformatics, 2011. 27(12): p. 1739–40.

37. Basisty, N., et al., A proteomic atlas of senescence-associated secretomes for aging biomarker development. PLOS Biology, 2020. 18(1): p. e3000599.

38. Avelar, R.A., et al., A multidimensional systems biology analysis of cellular senescence in aging and disease. Genome Biology, 2020. 21(1): p. 91.

39. Tate, J.G., et al., COSMIC: the Catalogue Of Somatic Mutations In Cancer. Nucleic Acids Research, 2018. 47(D1): p. D941–D947.

40. Zhao, M., J. Sun, and Z. Zhao, TSGene: a web resource for tumor suppressor genes. Nucleic Acids Res, 2013. 41(Database issue): p. D970–6.

41. Teschendorff, A.E. and T. Enver, Single-cell entropy for accurate estimation of differentiation potency from a cell’s transcriptome. Nature Communications, 2017. 8(1): p. 15599.

42. Cerami, E.G., et al., Pathway Commons, a web resource for biological pathway data. Nucleic Acids Research, 2010. 39(suppl_1): p. D685–D690.

43. Bader, G.D. and C.W. Hogue, An automated method for finding molecular complexes in large protein interaction networks. BMC Bioinformatics, 2003. 4: p. 2.

44. Wu, G. and R. Haw, Functional Interaction Network Construction and Analysis for Disease Discovery, in Protein Bioinformatics: From Protein Modifications and Networks to Proteomics, C.H. Wu, C.N. Arighi, and K.E. Ross, Editors. 2017, Springer New York: New York, NY. p. 235–253.

45. Street, K., et al., Slingshot: cell lineage and pseudotime inference for single-cell transcriptomics. BMC Genomics, 2018. 19(1): p. 477.

46. Dharmaratne, M., scShapes: A Statistical Framework for Modeling and Identifying Differential Distributions in Single-cell RNA-sequencing Data. R package version 1.0.0. 2021.

47. Fernandez-Rebollo, E., et al., Senescence-Associated Metabolomic Phenotype in Primary and iPSC-Derived Mesenchymal Stromal Cells. Stem Cell Reports, 2020. 14(2): p. 201–209.

48. Rohart, F., et al., A molecular classification of human mesenchymal stromal cells. PeerJ, 2016. 4: p. e1845.

49. de Witte, S.F.H., et al., Aging of bone marrow– and umbilical cord–derived mesenchymal stromal cells during expansion. Cytotherapy, 2017. 19(7): p. 798–807.

50. d’Adda di Fagagna, F., et al., A DNA damage checkpoint response in telomere-initiated senescence. Nature, 2003. 426(6963): p. 194–8.

51. Grasso, D., et al., Pivotal Role of the Chromatin Protein Nupr1 in Kras-Induced Senescence and Transformation. Scientific reports, 2015. 5: p. 17549–17549.

52. Nishimura, K., et al., Perturbation of Ribosome Biogenesis Drives Cells into Senescence through 5S RNP-Mediated p53 Activation. Cell Reports, 2015. 10(8): p. 1310–1323.

53. Yim, C.Y., et al., G0S2 Suppresses Oncogenic Transformation by Repressing a MYC-Regulated Transcriptional Program. Cancer Research, 2016. 76(5): p. 1204–1213.

54. Lafferty-Whyte, K., et al., TCEAL7 inhibition of c-Myc activity in alternative lengthening of telomeres regulates hTERT expression. Neoplasia (New York, N.Y.), 2010. 12(5): p. 405–414.

55. Royle, Nicola J., et al., The role of recombination in telomere length maintenance. Biochemical Society Transactions, 2009. 37(3): p. 589–595.

56. Guo, Y., et al., Senescence-associated tissue microenvironment promotes colon cancer formation through the secretory factor GDF15. Aging Cell, 2019. 18(6): p. e13013.

57. Guillon, J., et al., Regulation of senescence escape by TSP1 and CD47 following chemotherapy treatment. Cell Death & Disease, 2019. 10(3): p. 199.

58. Freund, A., et al., Inflammatory networks during cellular senescence: causes and consequences. Trends in Molecular Medicine, 2010. 16(5): p. 238–246.

59. Tominaga, K. and H.I. Suzuki, TGF-β Signaling in Cellular Senescence and Aging-Related Pathology. International journal of molecular sciences, 2019. 20(20): p. 5002.

60. Lyu, G., et al., Addendum: TGF-β signaling alters H4K20me3 status via miR-29 and contributes to cellular senescence and cardiac aging. Nature communications, 2018. 9(1): p. 4134–4134.

61. Kandhaya-Pillai, R., et al., SMAD4 mutations and cross-talk between TGF-β/IFNγ signaling accelerate rates of DNA damage and cellular senescence, resulting in a segmental progeroid syndrome—the Myhre syndrome. GeroScience, 2021. 43(3): p. 1481–1496.

62. Ren, J.-L., et al., Inflammatory signaling and cellular senescence. Cellular Signalling, 2009. 21(3): p. 378–383.

63. Lasry, A. and Y. Ben-Neriah, Senescence-associated inflammatory responses: aging and cancer perspectives. Trends in Immunology, 2015. 36(4): p. 217–228.

64. Day, P.J., et al., Crystal structure of human CDK4 in complex with a D-type cyclin. Proceedings of the National Academy of Sciences of the United States of America, 2009. 106(11): p. 4166–4170.

65. Sumi, T., et al., Survivin knockdown induces senescence in TTF-1-expressing, KRAS-mutant lung adenocarcinomas. International journal of oncology, 2018. 53(1): p. 33–46.

66. Chen, W.-S., et al., Ran-dependent TPX2 activation promotes acentrosomal microtubule nucleation in neurons. Scientific Reports, 2017. 7(1): p. 42297.

67. Hsu, C.-W., et al., Targeting TPX2 Suppresses the Tumorigenesis of Hepatocellular Carcinoma Cells Resulting in Arrested Mitotic Phase Progression and Increased Genomic Instability. Journal of Cancer, 2017. 8(8): p. 1378–1394.

68. Tang, H., et al., Single senescent cell sequencing reveals heterogeneity in senescent cells induced by telomere erosion. Protein & Cell, 2019. 10(5): p. 370–375.

69. Işildak, U., et al., Temporal changes in the gene expression heterogeneity during brain development and aging. Scientific Reports, 2020. 10(1): p. 4080.

70. Shmulevich, R. and V. Krizhanovsky, Cell Senescence, DNA Damage, and Metabolism. Antioxidants & Redox Signaling, 2021. 34(4): p. 324–334.

71. Gardner, J. and A. Ghorpade, Tissue inhibitor of metalloproteinase (TIMP)-1: the TIMPed balance of matrix metalloproteinases in the central nervous system. Journal of neuroscience research, 2003. 74(6): p. 801–806.

72. Dasgupta, J., et al., Reactive oxygen species control senescence-associated matrix metalloproteinase-1 through c-Jun-N-terminal kinase. Journal of cellular physiology, 2010. 225(1): p. 52–62.

73. Al-Mudares, F., et al., Role of Growth Differentiation Factor 15 in Lung Disease and Senescence: Potential Role Across the Lifespan. Frontiers in Medicine, 2020. 7(919).

74. Park, H., et al., GDF15 contributes to radiation-induced senescence through the ROS-mediated p16 pathway in human endothelial cells. Oncotarget, 2016. 7(9): p. 9634–9644.

75. Elzi, D.J., et al., Wnt antagonist SFRP1 functions as a secreted mediator of senescence. Molecular and cellular biology, 2012. 32(21): p. 4388–4399.

76. Su, Y., et al., S100A13 promotes senescence-associated secretory phenotype and cellular senescence via modulation of non-classical secretion of IL-1α. Aging, 2019. 11(2): p. 549–572.

77. Kadota, Y., et al., Gene expression of mesoderm-specific transcript is upregulated as preadipocytes differentiate to adipocytes in vitro. The Journal of Physiological Sciences, 2012. 62(5): p. 403–411.

78. Coulombe, P.A. and P. Wong, Cytoplasmic intermediate filaments revealed as dynamic and multipurpose scaffolds. Nature Cell Biology, 2004. 6(8): p. 699–706.

79. Victorelli, S. and J.F. Passos, Reactive Oxygen Species Detection in Senescent Cells. Methods Mol Biol, 2019. 1896: p. 21–29.

80. Puvvula, P.K., LncRNAs Regulatory Networks in Cellular Senescence. International journal of molecular sciences, 2019. 20(11): p. 2615.

81. Degirmenci, U. and S. Lei, Role of lncRNAs in Cellular Aging. Frontiers in Endocrinology, 2016. 7(151).

82. Zheng, D., et al., Cyclin-dependent kinase 3-mediated activating transcription factor 1 phosphorylation enhances cell transformation. Cancer research, 2008. 68(18): p. 7650–7660.

83. Arima, Y., et al., Down-regulation of nuclear protein ICBP90 by p53/p21Cip1/WAF1-dependent DNA-damage checkpoint signals contributes to cell cycle arrest at G1/S transition. Genes to Cells, 2004. 9(2): p. 131–142.

84. Bostick, M., et al., UHRF1 plays a role in maintaining DNA methylation in mammalian cells. Science, 2007. 317(5845): p. 1760–4.

85. Tien, A.L., et al., UHRF1 depletion causes a G2/M arrest, activation of DNA damage response and apoptosis. Biochem J, 2011. 435(1): p. 175–85.

86. Tousignant, K.D., et al., Therapy-induced lipid uptake and remodeling underpin ferroptosis hypersensitivity in prostate cancer. bioRxiv, 2020: p. 2020.01.08.899609.

87. Aguado, J., et al., Inhibition of DNA damage response at telomeres improves the detrimental phenotypes of Hutchinson–Gilford Progeria Syndrome. Nature Communications, 2019. 10(1): p. 4990.

88. Passos, J.F., et al., Mitochondrial dysfunction accounts for the stochastic heterogeneity in telomere-dependent senescence. PLoS Biol, 2007. 5(5): p. e110.

89. Kim, S.-J., et al., Mitochondrial peptides modulate mitochondrial function during cellular senescence. Aging, 2018. 10(6): p. 1239–1256.

90. Correia-Melo, C. and J.F. Passos, Mitochondria: Are they causal players in cellular senescence? Biochimica et Biophysica Acta (BBA) - Bioenergetics, 2015. 1847(11): p. 1373–1379.

91. Raj, A., et al., Stochastic mRNA Synthesis in Mammalian Cells. PLOS Biology, 2006. 4(10): p. e309.

92. Corrigan, A.M., et al., A continuum model of transcriptional bursting. eLife, 2016. 5: p. e13051.

