## Supplementary figures and tables for "Deconstructing replicative senescence heterogeneity of human mesenchymal stem cells at single cell resolution reveals therapeutically targetable senescent cell sub-populations": Sup_Figure_legends.docx

Figure S1. Expression of MSC-specific markers in the sc RNA-seq data prior to pre-processing. We inspected the scRNA-seq data to assess whether it met standard requirements for MSC characterisation. Traditionally, cells must express surface markers such as CD105 (ENG), CD73 (NT5E), and CD90 (THY1) and lack expression of several genes including CD14, CD19 and CD34. We observed that collectively, more that 79% of cells express (count > 0) either of the positive markers, and more than 99% of cells lack expression of (UMI = 0) negative markers.

Figure S2. Evaluating the expression of Rohart MSC signature (a) before and (b) after pre-processing the data. Out of the 16 MSC-specific genes in the Rohart MSC signature, 13 genes were identified in our dataset prior to filtering for lowly expressed genes; and four genes remained after filtering (*i.e.* removing genes with less than 10% expression in all cells). More than 70% of the cells expressed ITGA11 and 41.14% of cells expressed PRRX1. This was followed by GDF5 and MBD2 with 19.36% and 14.78% of cells respectively

Figure S3. Validating the senescence phenotype through senescence specific markers and phenotypic features, for individual replicates.

Figure S4. Telomere length and DNA damage analysis through replicative senesce in esMSCs in replicates 2 and 3. (a) Normalised telomere length across all time-points. (b) Telomere Dysfunction Induced Foci (TIF) analysis (R2: p-value = ns; R3: p-value < 0.001). The violin plot shows the number of cells that indicated DNA double strand break at the telomeres at each time-point. There is no significant difference between the three time-points (one-way ANOVA, p-value > 0.05 for all pairwise comparisons) (c) Genome wide DNA double strand break. The stack bar shows the percentage of cells that stained positive for γH2AX, genome wide. The number of cells with DNA damage increases from T0 to T2, however the extent of change is not statistically significant. (d) DNA double-strand break analysis at the telomeres. The stack bar shows the percentage of cells that stained positive for γH2AX, at the telomeres. Similar to the genome-wide analysis, we observe a gradual increase in number of cells from T0 to T2 but the difference in proportions are not statistically significant. *ns= non-significant.

Figure S5. SenezRed staining show that there is no significant difference between different cells at different timepoints and replicates.

Figure S6. Correlation analysis between senescence specific markers and phenotypic features. Pearson correlation was computed for each marker intensity (samples are pooled from replicate 2 and 3).

Figure S7. UMAP of cells, color coded by cluster and split by timepoint. Clusters 2, 6 and parts of cluster 3 comprise of T0 cells. Cluster 5 and 0 comprise of T1 cells, and proportion of cluster 3, 5 and 0, and majority of cluster 1 and 4 comprise of T2 cells.

Figure S8. Normalised gene expression of top five marker genes for each (a) cluster and (b) time-point.

Figure S9. Differential expression analysis of T1 cells between different clusters. According to pseudotime trajectory analysis, cluster 0 is the main point of divergence between the two trajectories. Thus, we chose T1 cells in cluster 0 (shown as red) as the base of comparison with T1 cells in other clusters (shown in blue and grey).

Figure S10. Differential expression analysis of senescence escapees (in blue) with (a) Proliferative esMSCs subtype-3 (other cells in cluster 3), and (b) other senescent cells (other T2 cells). Top panel: UMAP highlight cells in the comparison. Bottom panel: Significant DE genes (LogFC x > |0.25|, Bonferroni adjusted p-value<0.001). (c) Overlap between differentially expressed genes of senescence escapees with proliferative esMSC subtype-3 (other cells in cluster 3), and other senescent cells (other T2 cells). The Venn diagram shows the overlap between differentially expressed genes of senescence escapees with proliferative esMSCs subtype-3 (other cells in cluster 3), and other senescent cells (other T2 cells).

Figure S11. Functional interaction (FI) network analysis at each time-point. Nodes represent the genes in the PPI network (extracted via SCENT package) and edges represent the spearman-correlation computed from gene expression data. Highly connected modules were obtained via MCODE app in Cytoscape. Red and green edge colors represent positive and negative correlations, respectively. Edge types were extracted from ReactomeFIViz app in Cytoscape which has access to the Reactome pathway database. They show the mode of regulatory interaction: "->" for activating/catalyzing, "-|" for inhibition, "-" for FIs extracted from complexes or inputs, and "---" for predicted FIs.

Figure S12. Normalised expression of mtDNA genes during senescence. Violin plots of each mtDNA gene across three different time points. *MT-ATP8, MT-ND4L and NT-ND6 are not shown due to an overall low UMI count after pre-processing and normalisation.
