## Supplementary figures and images for "Deconstructing replicative senescence heterogeneity of human mesenchymal stem cells at single cell resolution reveals therapeutically targetable senescent cell sub-populations"

### FigSup1_msc_markers.jpg

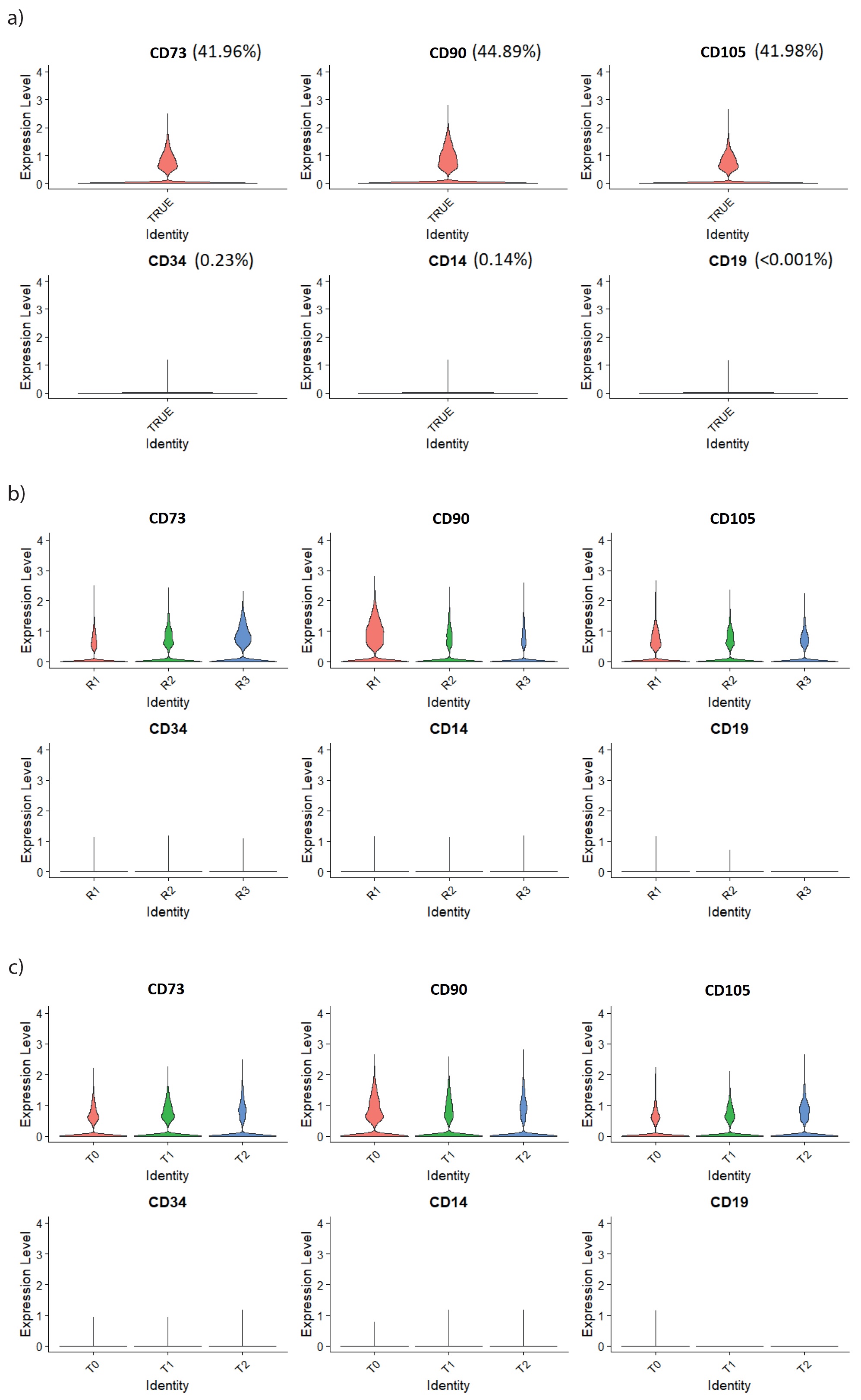

### FigSup2_rohart_markers.jpg

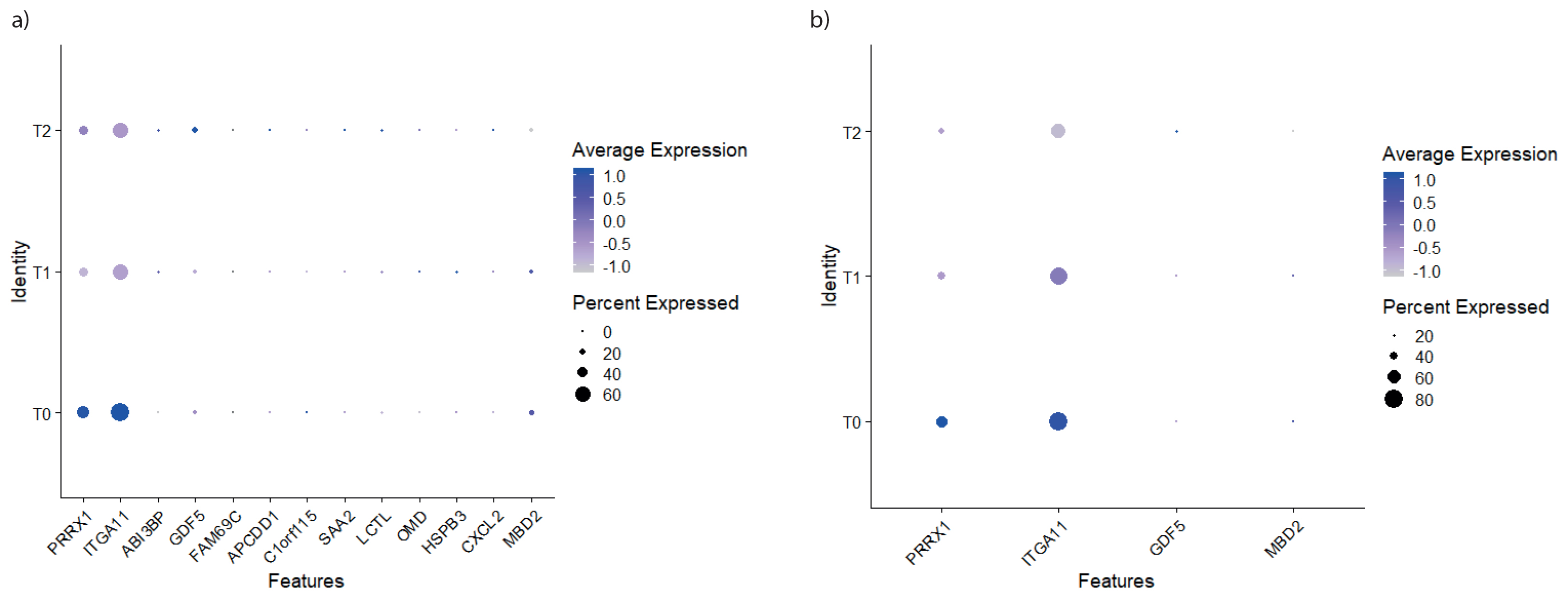

### FigSup3_Violin plot for all scaled intensity per replicate.png

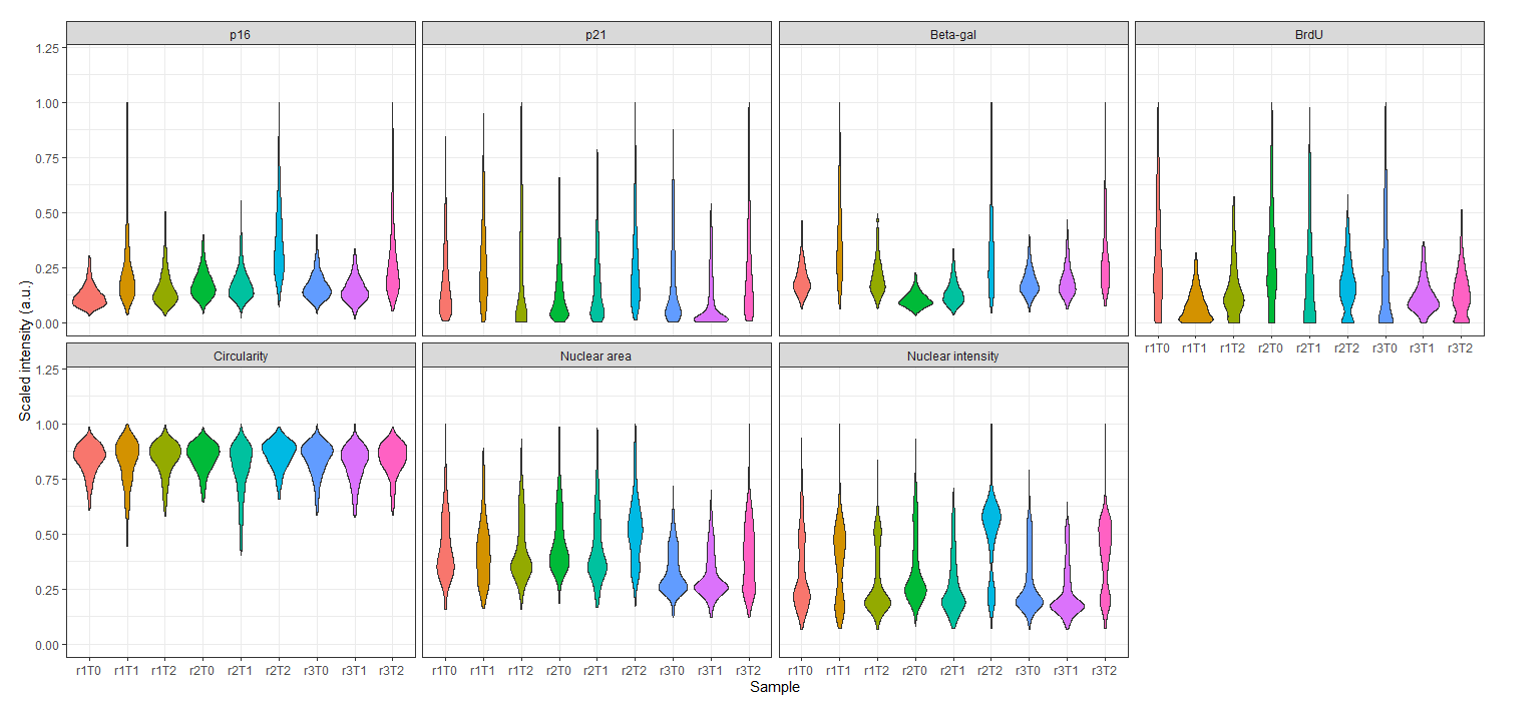

### FigSup4_TIF_DNAdamage.jpg

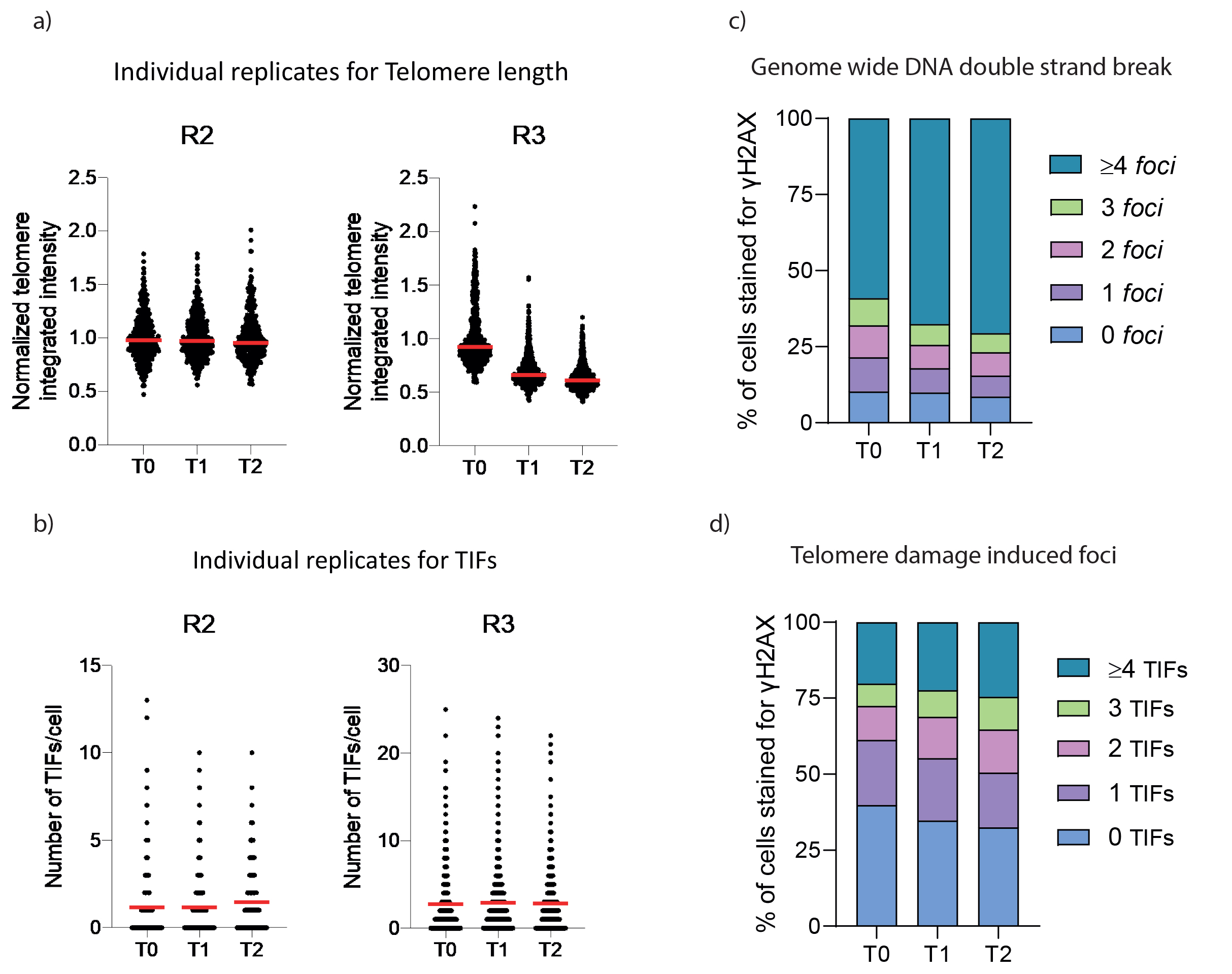

### FigSup5_MSC SenZ analysis.jpg

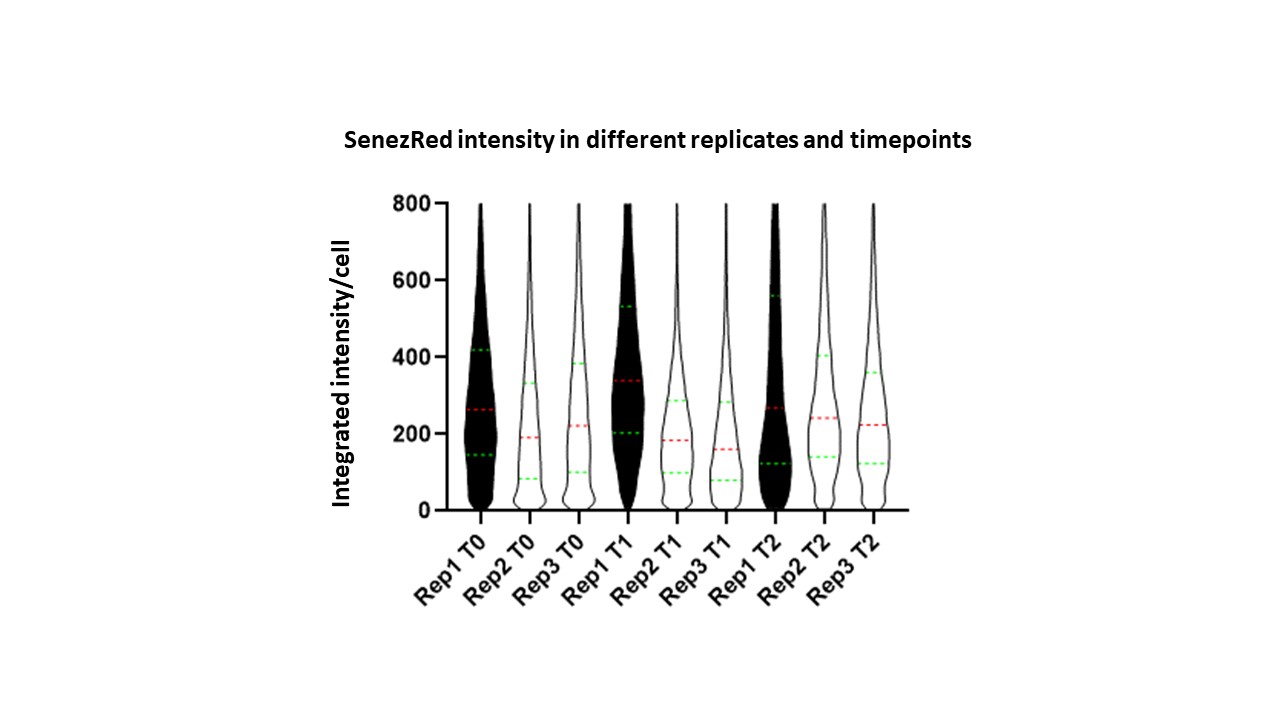

### FigSup6_Correlations of markers across time point -1.png

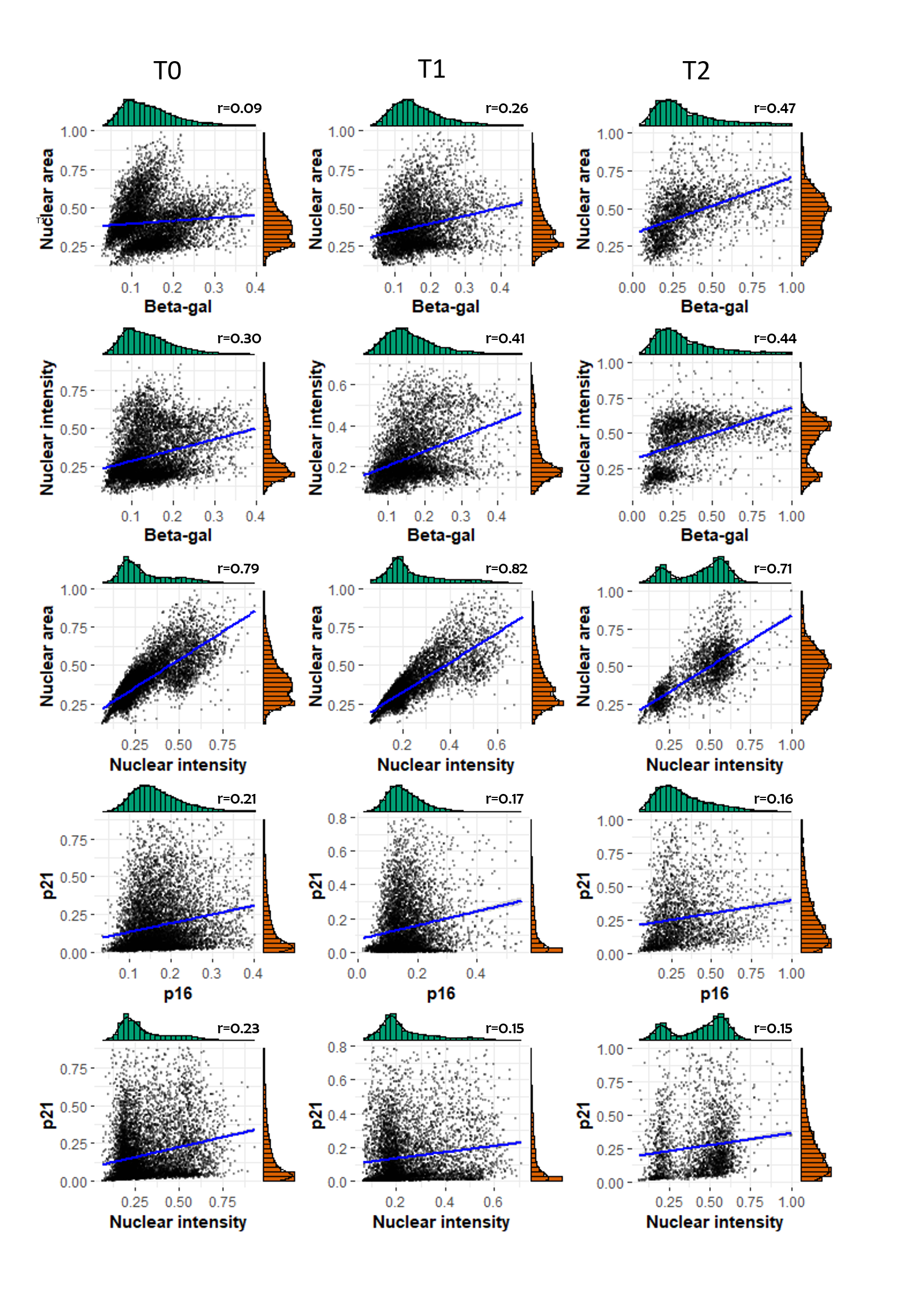

### FigSup6_Correlations of markers across time point.png

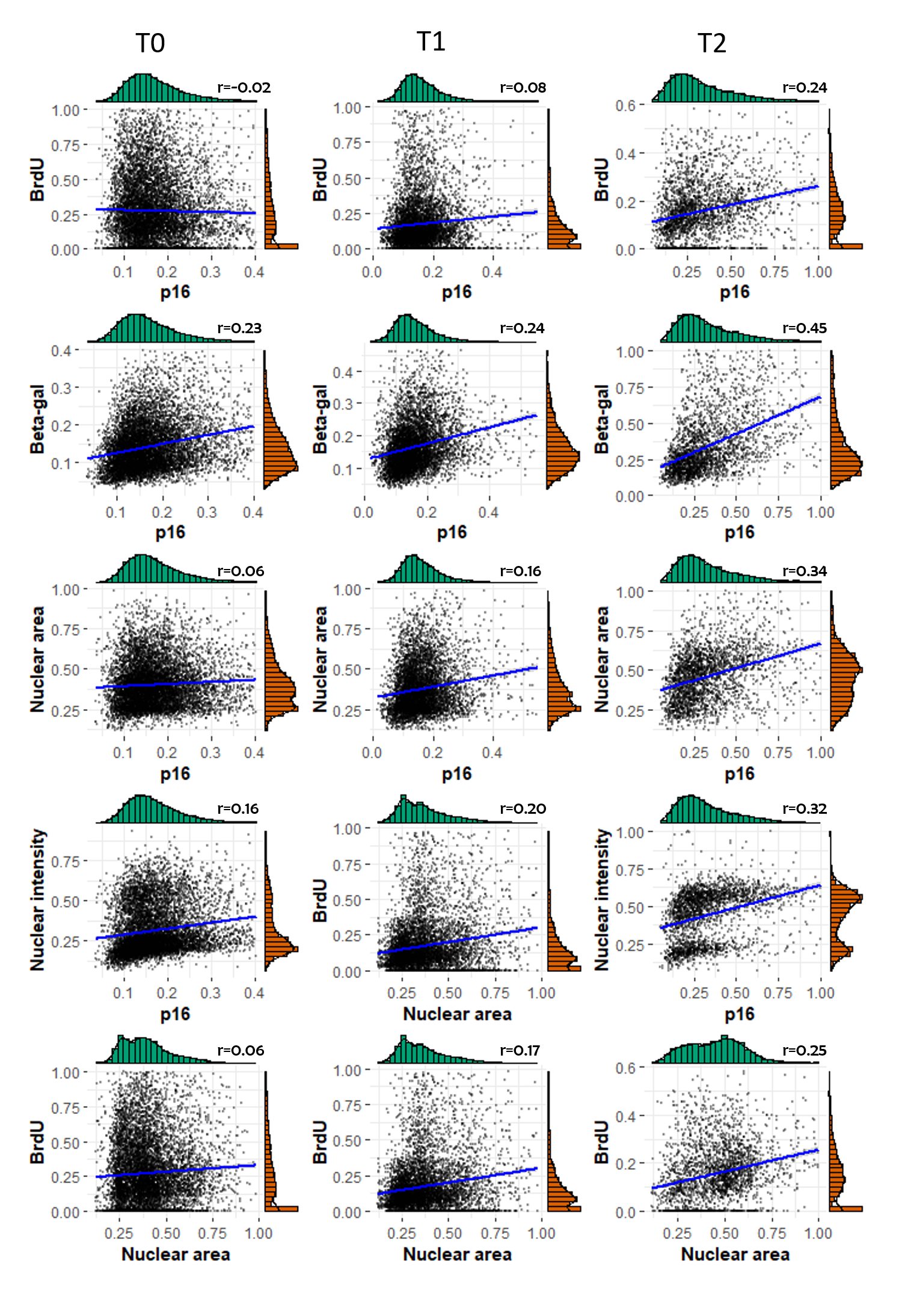

### FigSup7_umap_integrate_sct_clust&group_10p.png

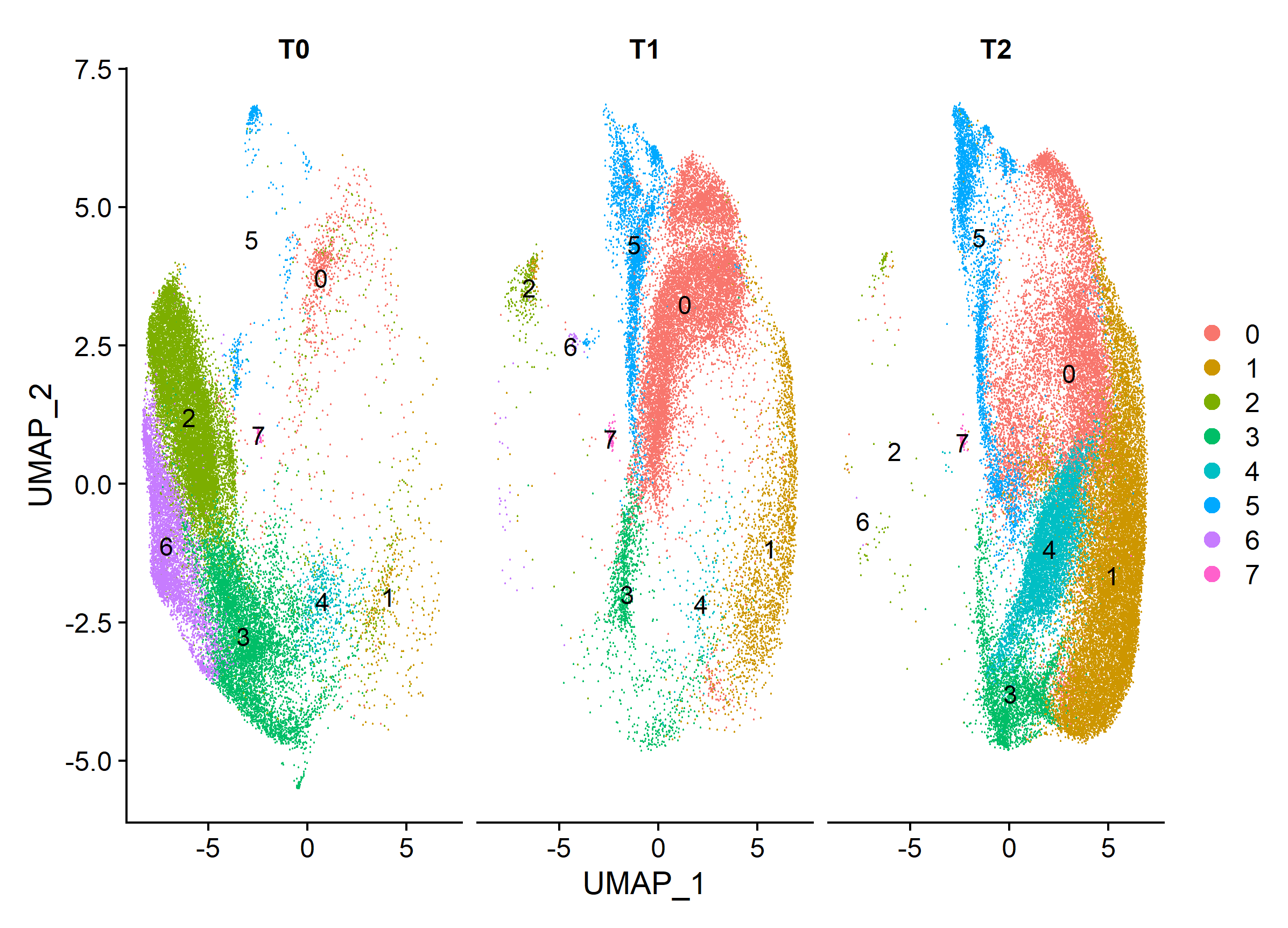

### FigSup8_top5DE.jpg

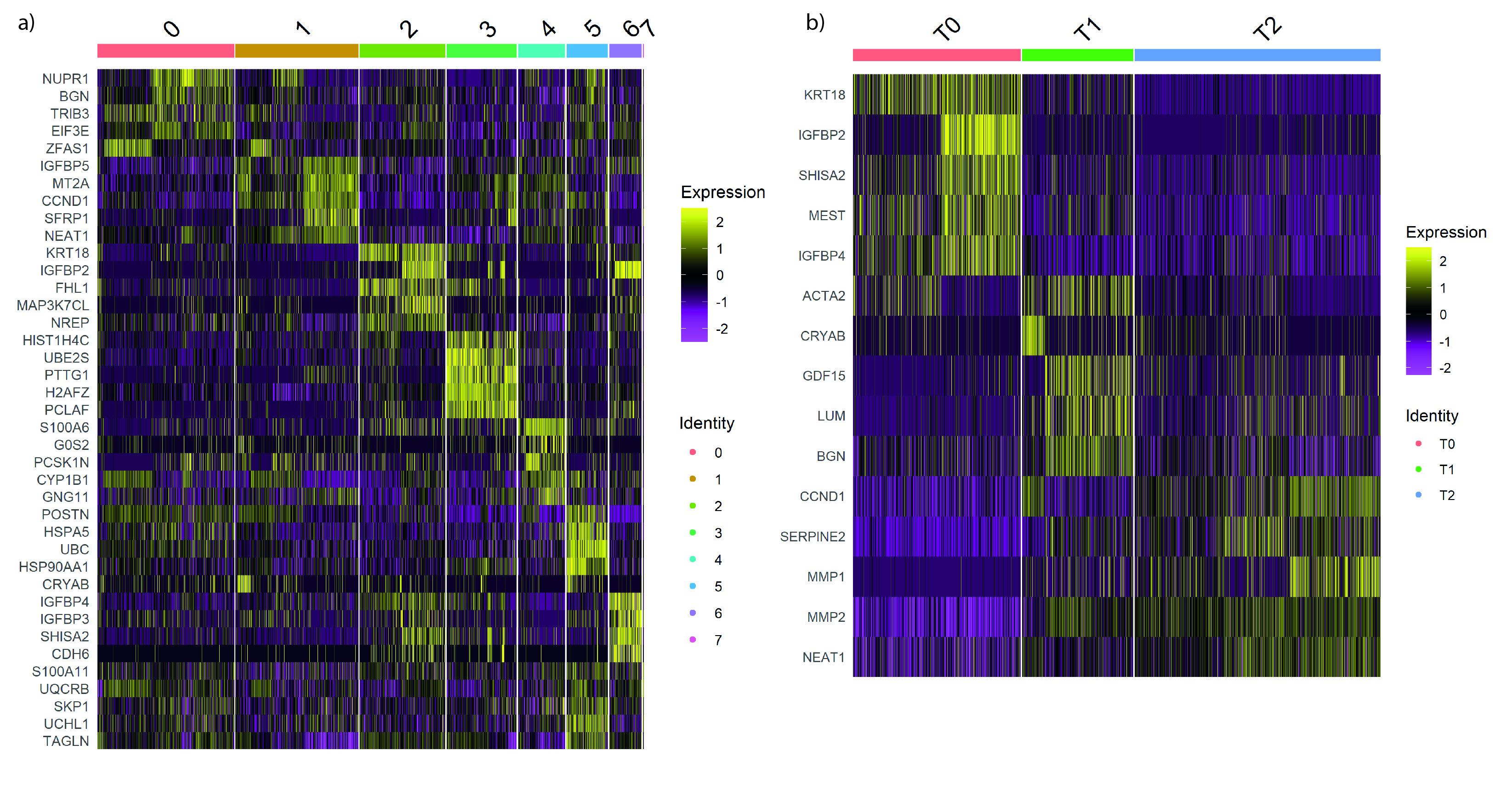

### FigSup9_T1_comparison.jpg

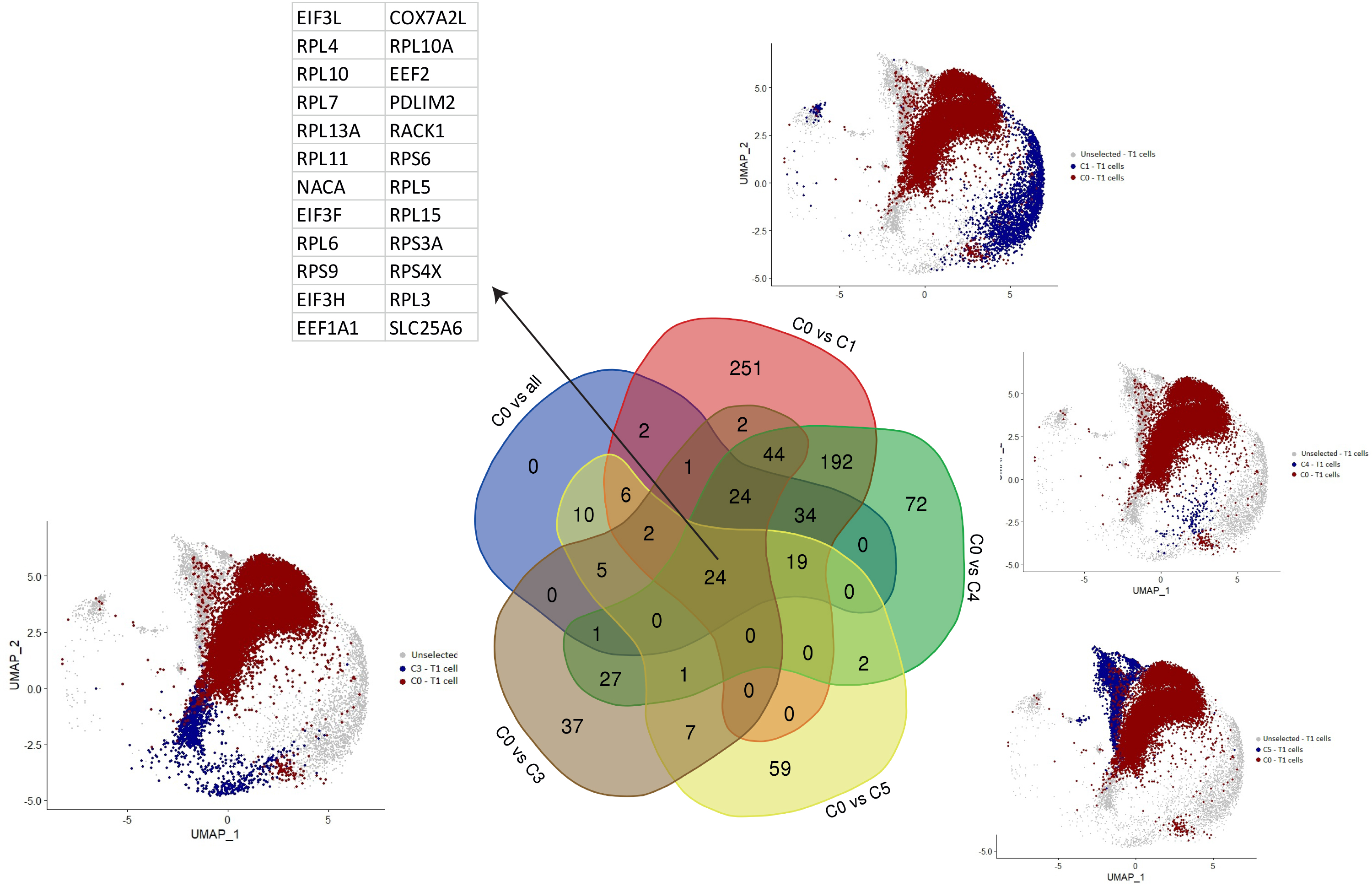

### FigSup10_DE_escapees.jpg

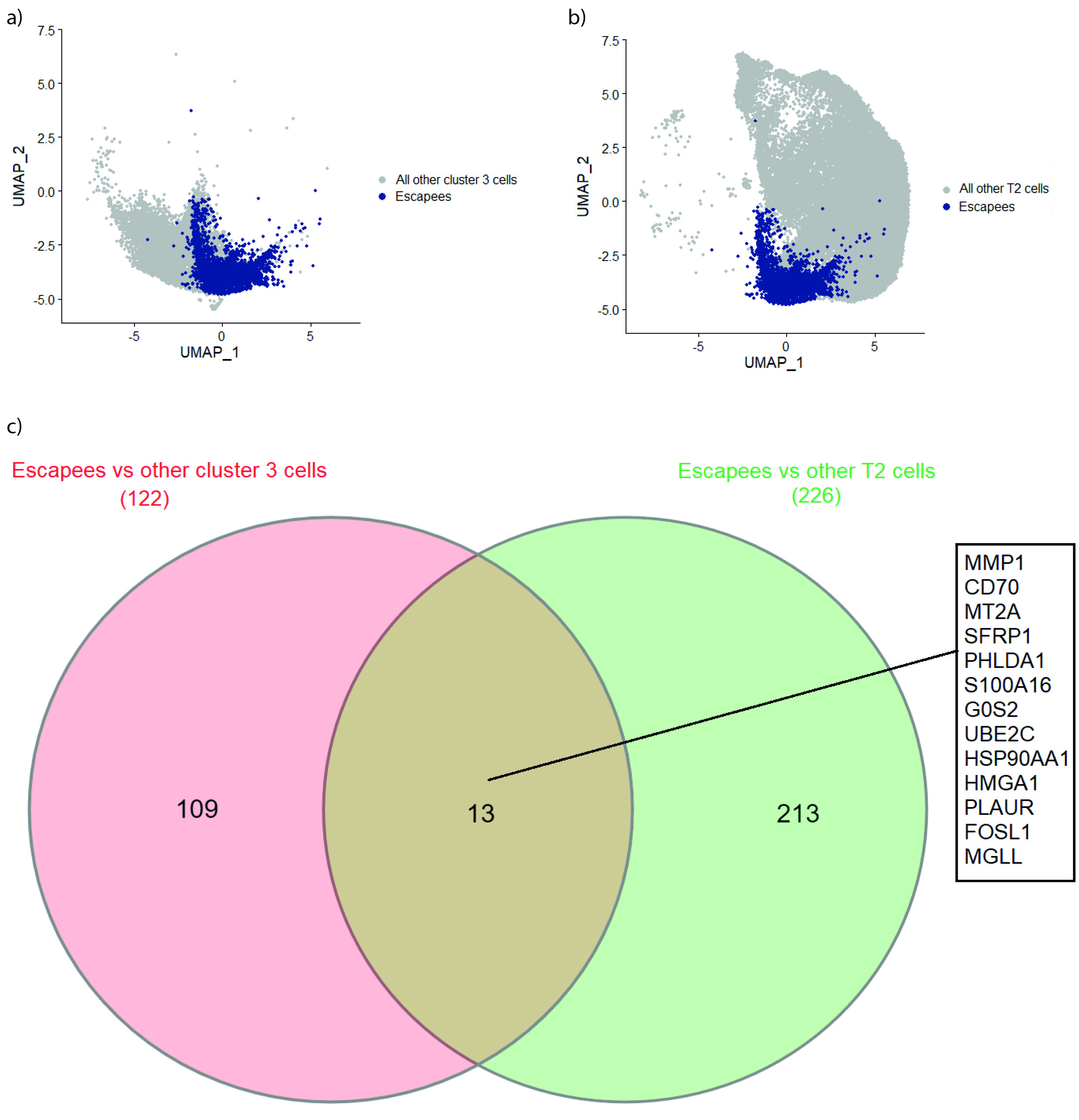

### FigSup11_ppi_scent1.jpg

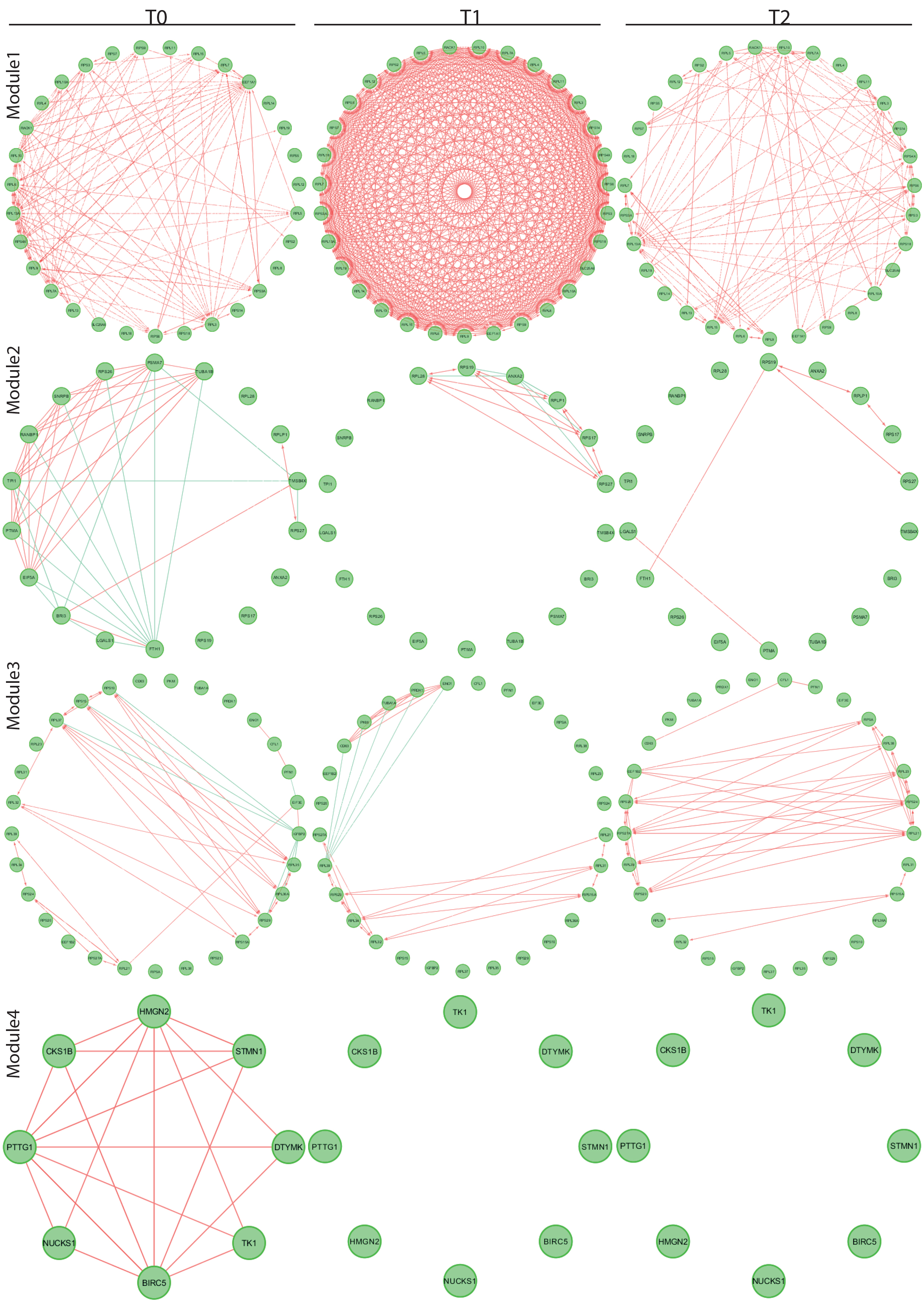

### FigSup11_ppi_scent2.jpg

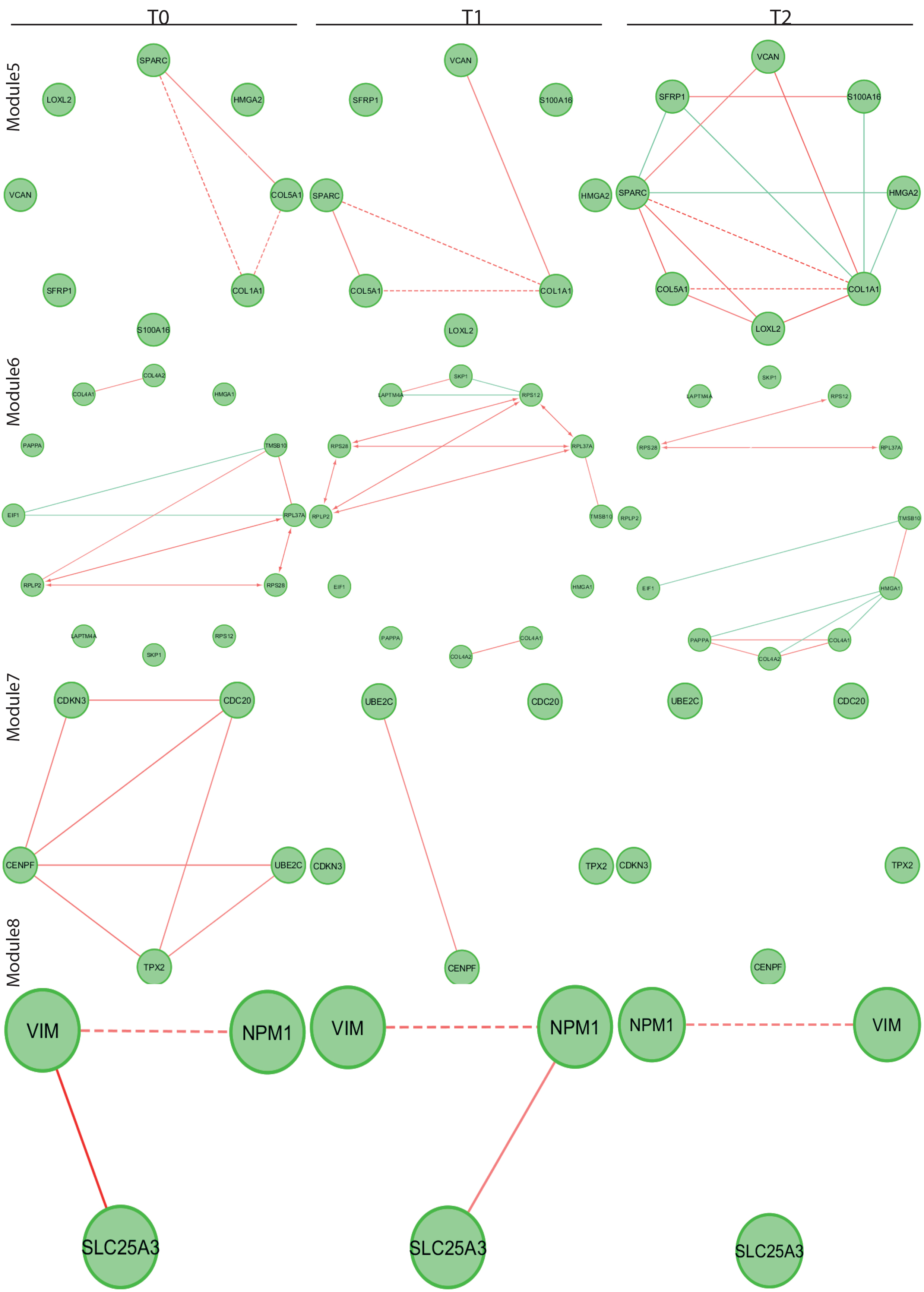

### FigSup12_mtDNA_violinPlots.png

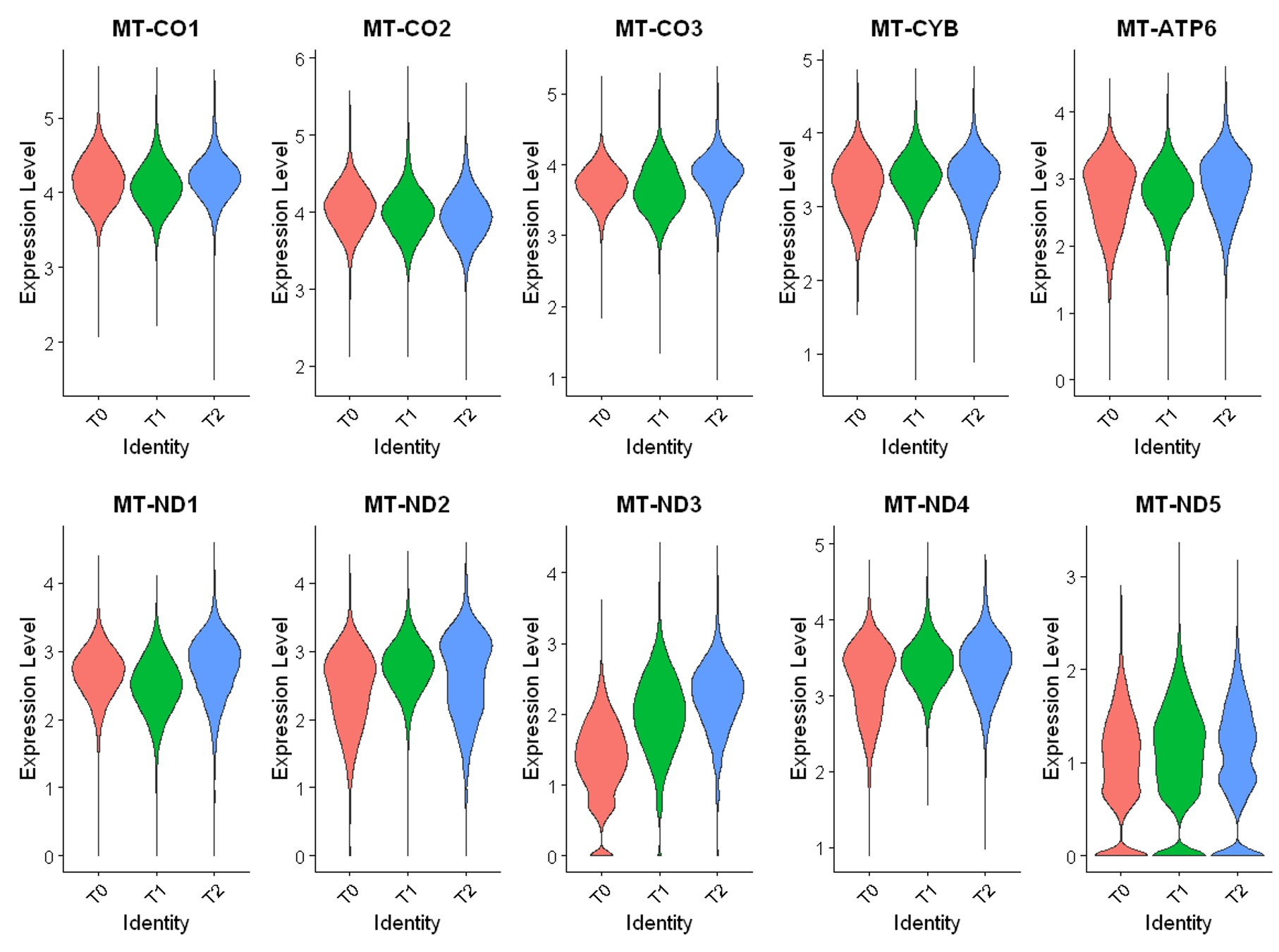
